# *U6* promoters in *Ulva prolifera* enable dual small-RNA/mRNA transcription and vector-based CRISPR-Cas9 genome editing

**DOI:** 10.64898/2026.10.23.761446

**Authors:** Qingyun Hou, Zhenghua Wang, Jin Zhao, Peng Jiang

## Abstract

For establishing vector-based gene editing in the green-tide-forming alga *Ulva prolifera*, development of endogenous RNA polymerase III (Pol III) promoters is a prerequisite to drive single-guide RNA (sgRNA) transcription. To meet the need, the promoters of *U6* snRNA genes in *U. prolifera* (*UpU6*) were identified, cloned, and functionally characterized. Results showed that three *UpU6* copies shared identical sequences, with divergent 5’ upstream sequences containing a positionally conserved TATA box but lacking the USE element typical of seed plants. All three *UpU6* promoters were shown to be capable of driving not only small RNA transcription but also protein-coding gene expression in *U. prolifera*, exhibiting the dual transcriptional activity. Unexpectedly, all three *UpU6* promoters also supported *GUS* reporter gene expression in *Agrobacterium*, representing the first demonstration of *U6* promoter activity in a prokaryotic system. The results of the gene-editing experiment showed that, all three *UpU6* promoters supported vector-based CRISPR-Cas9 targeting of the *APT* gene, yielding mutant strains. The mutation-positive fraction among resistant thalli was 92.3% (48/52), and the editing frequency calculated from the estimated starting cell number averaged 3.26 × 10⁻⁶ across the three *UpU6* promoters and did not differ significantly among them. In the mutants, high-frequency occurrences of base substitutions (45.5%) and insertions of long Mutator-like element (MULE) (18.2%) were detected, highlighting a potential association between endogenous DNA transposon insertion and CRISPR-Cas9-mediated gene editing. These findings expanded understanding of eukaryotic *U6* promoter functionality and enrich the toolkit for vector-based CRISPR-Cas9 genome editing in *U. prolifera*.

## 1. Introduction

The genus *Ulva*, a group of green algae, is widely distributed in global coastal marine environments. As a typical “opportunistic” seaweed, it can rapidly respond to excessive nutrient conditions in seawater, resulting in substantial biomass accumulation and the formation of green tides [1]. Since 2007, the Yellow Sea green tide (YSGT) dominated by *Ulva prolifera* has occurred annually, with the maximum distribution area exceeding 60000 km² and total biomass reaching several million tons [2,3]. The YSGT not only causes oxygen depletion in water bodies and releases harmful substances, leading to extensive mortality of marine organisms, but also inflicts severe economic losses [4,5]. Genetic analyses have demonstrated that the YSGT is caused by *U. prolifera* with genetic specificity [6,7], highlighting the necessity of elucidating the biological mechanisms underlying its massive blooms. To date, the genome of *U. prolifera* has been sequenced [8,9], and recent studies integrating multi-omics approaches with physiological and biochemical analyses have provided important insights into its environmental adaptation mechanisms [10–12]. Currently, the establishment of efficient gene-editing technologies for *U. prolifera* is urgently needed to enable functional validation of candidate genes potentially crucial to algal bloom formation.

The CRISPR-Cas9 system is currently the most widely adopted genome-editing technology, which relies on the Cas9 nuclease and a single-guide RNA (sgRNA) to induce targeted DNA double-strand breaks (DSBs) [13,14]. Depending on the delivery method for Cas9 and sgRNA, the construction of CRISPR-Cas9 systems primarily employs two strategies: ribonucleoprotein (RNP)-based delivery and vector-based delivery. The former directly delivers a complex of Cas9 protein and *in vitro*-synthesized sgRNA, whereas the latter introduces a plasmid carrying the genes encoding Cas9 and sgRNA. To date, effective genetic transformation methods have been established in *Ulva*, including polyethylene glycol (PEG)-mediated transformation [15,16] and biolistic delivery via gene gun [17–19]. Based on this, gene knockout and knock-in have been successfully achieved in *U. prolifera* using RNP delivery [20–22]. Compared with the RNP approach, vector-based delivery offers distinct advantages in enabling efficient screening of mutants, reducing costs and operational complexity, and facilitating complex editing strategies such as multiplex genome editing and base editors. Currently, a vector-based approach has been established in *U. compressa*, enabling duplex genome editing, and it has been proposed that further optimization of different genetic elements within the vector architecture could still improve editing efficiency [23]. However, this method has not yet been attempted in the green-tide alga *U. prolifera*.

In vector-based CRISPR-Cas9 systems, Cas9 gene transcription is driven by an RNA polymerase II (Pol II) promoter, whereas sgRNA transcription relies on an RNA polymerase III (Pol III) promoter [24]. As an RNA-guided DNA endonuclease system, editing efficiency is largely determined by sgRNA activity and the abundance of Cas9–sgRNA complexes [25,26]. Consequently, promoter selection is critical for optimizing the expression of these components. In eukaryotes, U6 small nuclear RNA (snRNA) is one of the most abundant snRNAs involved in pre-mRNA splicing and is transcribed by Pol III [27,28]. Owing to its conserved transcription start site (TSS) and termination signals, the *U6* promoter enables precise sgRNA transcription, thereby enhancing targeting specificity and reducing off-target effects [29,30]. As a result, it has become the most widely used Pol III promoter to date [29,31], and attempts have been made in *U. compressa* [23].

Multiple studies have shown that using an endogenous *U6* promoter within a species can significantly improve gene editing efficiency compared to exogenous *U6* promoters [32–35]. Furthermore, the *U6* snRNA gene is typically present in multiple copies in eukaryotes [36,37]. *U6* promoter sequences from different copies exhibit substantial divergence, with transcriptional activity differences reaching up to 10-fold [38–40]. Therefore, to construct an efficient CRISPR-Cas9 gene editing system in *U. prolifera*, it is necessary to identify, clone, and evaluate the transcriptional activities of all endogenous *U6* promoters. This need is particularly pronounced in multiplex genome editing systems, where multiple highly active *U6* promoters are essential [40–42].

In *U. prolifera*, exogenous gene transient and stable expression systems have been established respectively [18,19], providing a critical tool for evaluating the efficacy of *U6* promoters in small RNA transcription and gene editing. Beyond this, the potential functional flexibility of *U6* promoters needs to be specifically examined. In vertebrates, *U6* promoters have been shown to recruit not only Pol III for snRNA transcription but also Pol II for mRNA transcription [43,44]. This dual polymerase activity may influence the transcriptional balance between desired sgRNAs and unintended long transcripts, thereby affecting sgRNA expression levels and gene editing efficiency under specific conditions [44]. In various plants, *U6* promoters have also been demonstrated to drive reporter gene expression [30,33,40,42,45,46], suggesting that such dual activity may be also prevalent in plants. In the early divergent lineage of the *Ulva* genus, although the *U6* promoter from *U. compressa* has been applied in gene editing [23], it remains unclear whether the *U6* promoter possesses flexible transcriptional capabilities. Furthermore, studies have suggested that Pol III promoter activity influences sgRNA expression levels, which may in turn affect CRISPR editing efficiency and DNA repair outcomes [25,29,47]. This is also within the scope of this research.

In this study, all *U6* snRNA gene copies and their upstream regulatory sequences in the *U. prolifera* genome were identified and cloned. Through vector construction combined with established genetic manipulation systems, the activity of each *U6* promoter was systematically evaluated in driving transcription of both mRNA and small RNA in *U. prolifera*, as well as in supporting *GUS* gene expression in the prokaryotic host. Furthermore, using a vector-based gene editing approach, the editing outcomes associated with different *U6* promoter copies were evaluated. These findings would not only expand the understanding of the functional flexibility of *U6* promoters, but also provide a toolkit of regulatory elements for the CRISPR-Cas9 genome editing platform in *U. prolifera*.

## 2. Materials and methods

### 2.1 Algal materials, bacterial strains and Arabidopsis thaliana

*Ulva prolifera* O.F.Müller strain S096 was maintained as a unialgal culture in our laboratory. This strain was originally isolated from floating green macroalgae offshore Qingdao during the YSGT event in 2007 [48]. Species identity, ecotype, and life-cycle generation were verified using multiple genetic markers following established protocols [6,49–51], confirming its designation as a sporophyte of *U. prolifera* floating ecotype. For vegetative growth and biomass accumulation, algal thalli were cultured in Von Stosch’s Enriched (VSE) medium with weekly renewal, under controlled conditions of 16°C, a 12 h:12 h light:dark photoperiod, and a photosynthetic photon flux density of approximately 60 µmol photons m^-2^ • s^-1^. Chemically competent *Escherichia coli* TOP10 cells were purchased from Tolo Biotech. Co., Ltd. (Shanghai, China). *Agrobacterium tumefaciens* LBA4404 chemically competent cells were obtained from Shanghai Angyu Biotechnology Co., Ltd. (Shanghai, China). Seeds of *Arabidopsis thaliana* Columbia ecotype were kindly provided by Prof. Jingshan Wang at Qingdao Agricultural University.

### 2.2 Cloning and sequence analysis of UpU6 genes with 5’ upstream regions

Genomic DNA was extracted from *U. prolifera* using a Plant Genomic DNA Extraction Kit (Tiangen Biotech Co., Ltd., Beijing, China) following the manufacturer’s protocol. DNA integrity was assessed by electrophoresis on a 1% (w/v) TAE agarose gel. To identify *U6* snRNA genes of *U. prolifera* (*UpU6*), the complete sequence of *U6* snRNA gene of *Chlamydomonas reinhardtii* (*CrU6*; GenBank: X71486.1) was used as a query for BLASTN searches against three available *U. prolifera* genome assemblies: two from floating ecotypes associated with the YSGT (GenBank: JALAAN000000000.1 and SDUY00000000.1) [8] and one from an attached population in Hiroshima, Japan (GenBank: BRCE00000000.1) [9].

For each highly homologous genomic copy identified, a primer pair was designed using Primer Premier 5 (Table S1) to amplify a fragment spanning the putative *UpU6* gene and approximately 1.0 kb of its 5’ upstream region. PCR amplification was performed using ApexHF HS DNA Polymerase FS Master Mix (AG12202, Accurate Biology Co., Ltd., Changsha, China) and *U. prolifera* genomic DNA as template, with a T960 thermal cycler (Heal Force, Shanghai, China). The cycling conditions consisted of an initial denaturation at 94°C for 30 s, followed by 35 cycles of 98°C for 10 s, 55°C for 15 s, and 72°C for 15 s. PCR products were purified using the E.Z.N.A.® Gel Extraction Kit (Omega Bio-tek, Norcross, GA, USA). A 3’ A-overhang was added to the blunt-ended products by incubation at 72°C for 20 min with 2× Taq Master Mix (Novoprotein Scientific Inc., Suzhou, China) and High Pure dNTPs (TransGen Biotech Co., Ltd., Beijing, China). The A-tailed fragments were ligated into the pGEM-T vector (Promega, Madison, WI, USA), which contains a complementary T-overhang, and the resulting plasmids were transformed into *E. coli* TOP10 competent cells. Plasmid DNA was extracted and submitted to Sangon Biotech Co., Ltd. (Shanghai, China) for Sanger sequencing.

**Table S1.**
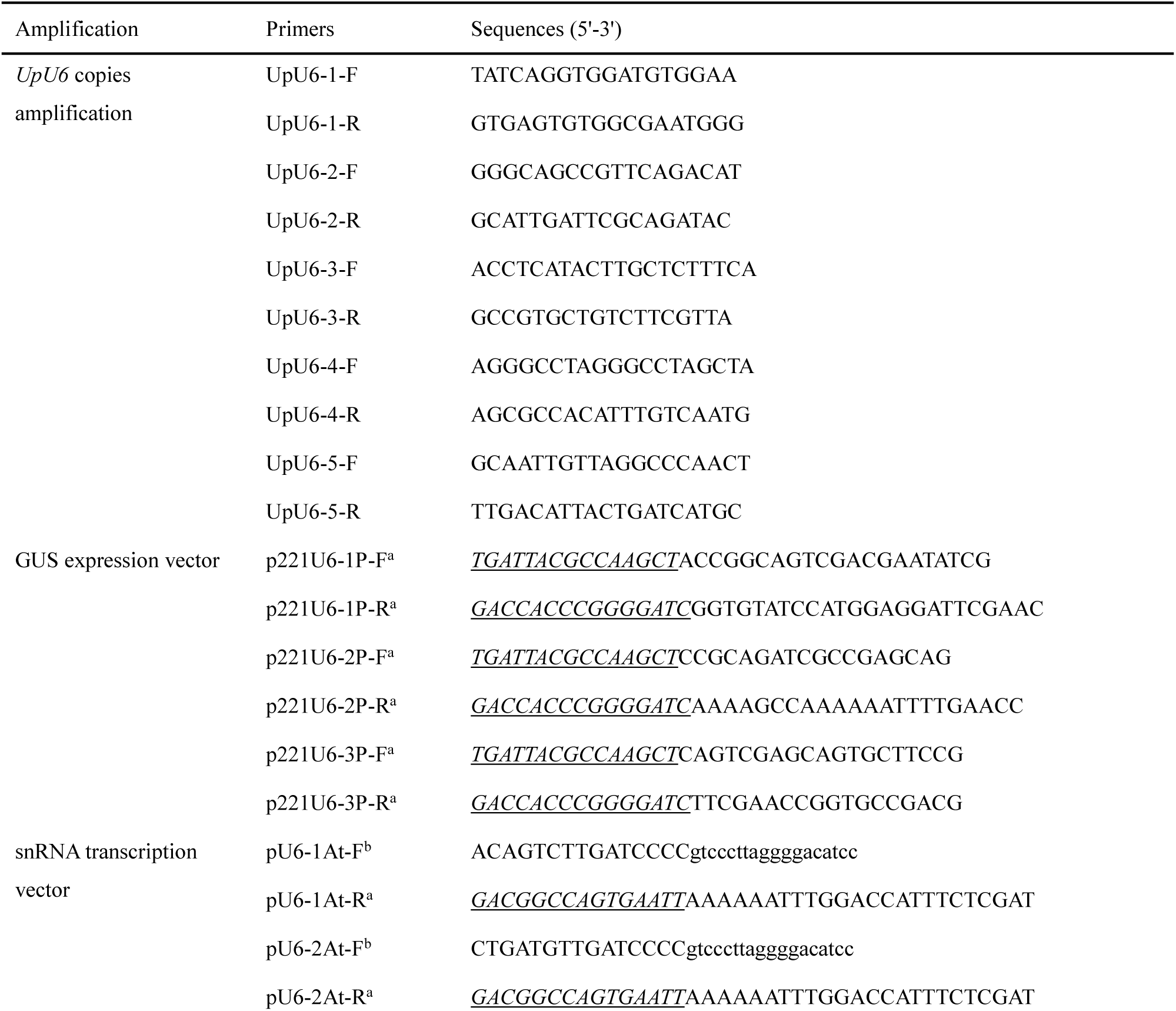
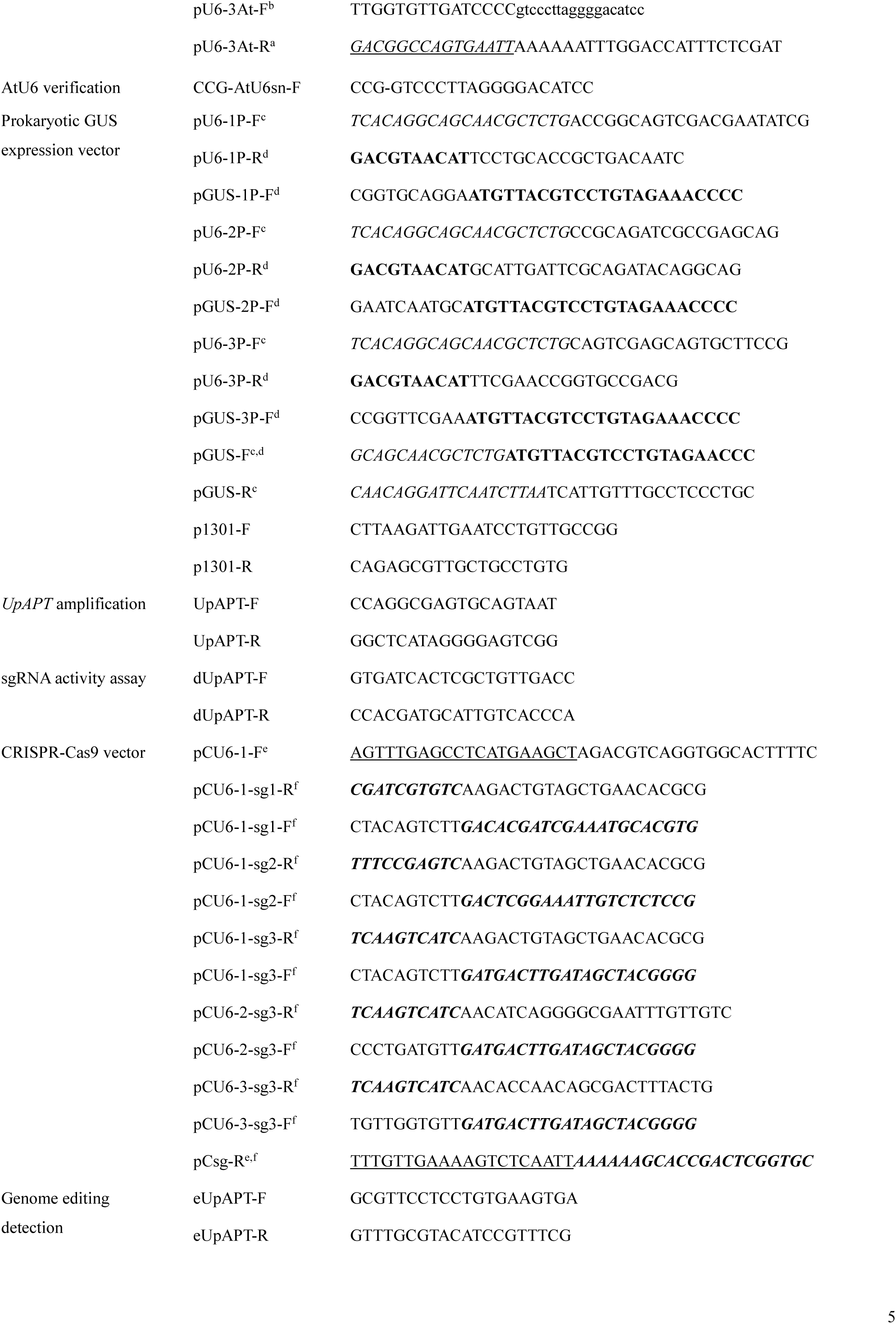
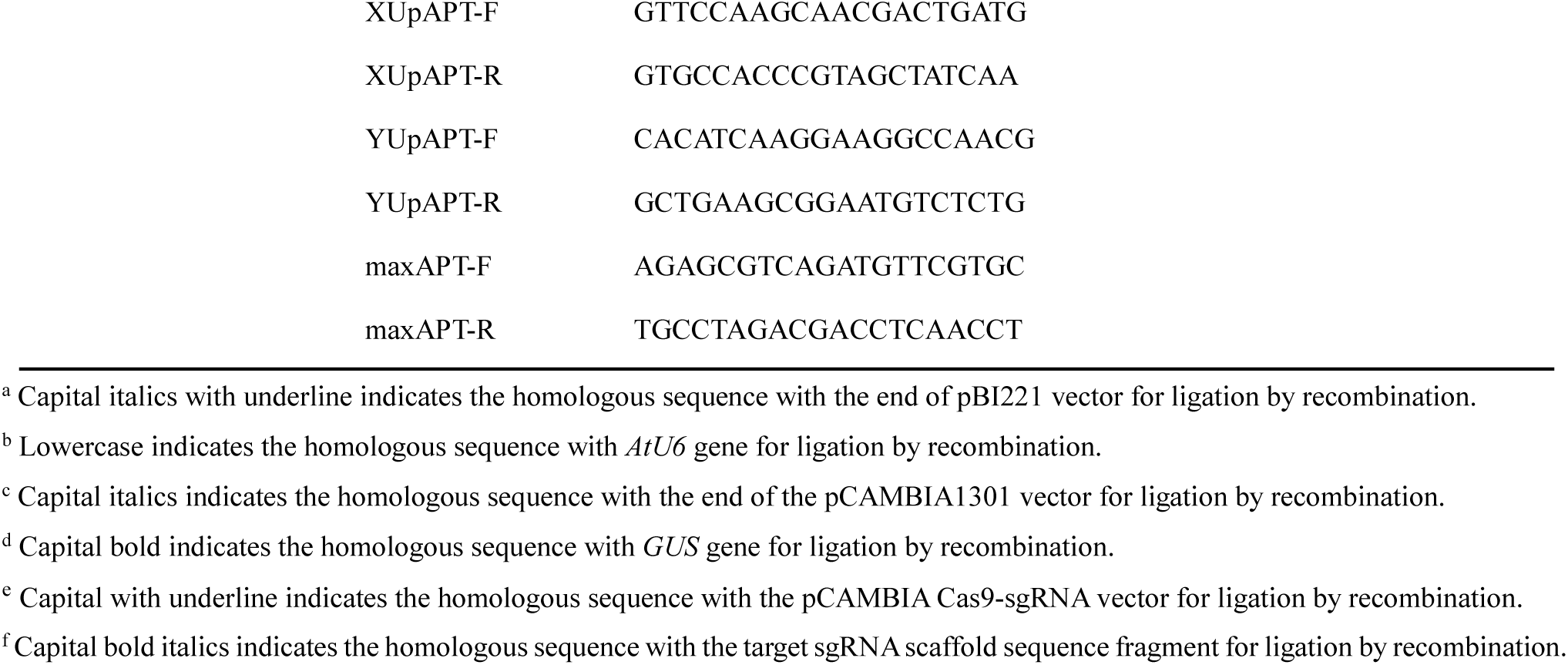
All primers used in this research.

To determine the complete sequence of each *UpU6* gene copy and to identify conserved transcriptional regulatory elements within the 5’ upstream regions, all obtained sequences were aligned using DNAMAN 6.0 (Lynnon BioSoft, San Ramon, CA, USA) together with orthologous sequences from representative plant species retrieved from the GenBank database. Potential *cis*-regulatory elements within the 5’ upstream region of each *UpU6* copy were further analyzed using PlantCARE (http://bioinformatics.psb.ugent.be/webtools/plantcare/html/) [52].

### 2.3 Construction of vectors

To evaluate the function of *UpU6* promoters in driving mRNA transcription, pBI221 (Clontech, Mountain View, CA, USA) was used as the backbone vector to construct a series of promoter-probe vectors. The native CaMV35S promoter was excised from pBI221 using *Hin*dIII and *Bam*HI (New England Biolabs, Ipswich, MA, USA) and replaced with approximately 1.0 kb of the 5’ upstream region from each *UpU6* genomic copy respectively. PCR products were purified using the E.Z.N.A.® Gel Extraction Kit, and cloned using the Seamless Assembly Cloning Kit (CloneSmarter, Fort Bend, TX, USA).

In addition, to further verify whether each *UpU6* promoter can drive *in vivo* transcription of snRNA in *U. proifera*, the entire CaMV35S promoter-GUS-nos terminator cassette was replaced with a fusion fragment comprising approximately 300 bp of the 5’ upstream region from one *UpU6* copy and the complete heterologous *U6* snRNA gene from *A. thaliana* (*AtU6*).

Furthermore, to evaluate the function of *UpU6* promoters in driving mRNA transcription in prokaryotes, expression vectors were constructed by fusing the *GUS* reporter gene to each *UpU6* promoter. Given that the endogenous GUS background of *E. coli* can interfere with detection [53], *A. tummefaciens* LBA4404 was selected as the host instead. The binary vector pCAMBIA1301 (Tiandz, Beijing, China) served as the backbone for vector construction, with the original prokaryotic promoter removed to avoid potential interference. Since the *GUS* gene in pCAMBIA1301 is inactivated by an intron in *Agrobacterium*, the intron-free *GUS* gene was amplified from pBI221 and used to replace the original. The approximately 300-bp 5′ upstream region of each *UpU6* copy was assembled separately with the intron-free *GUS* reporter gene and the modified vector backbone using the Seamless Assembly Cloning Kit to generate the promoter probe vectors. A promoterless negative control was constructed using the same modified vector backbone and intron-free *GUS* reporter gene, but without the *UpU6* promoter region. Finally, CRISPR-Cas9 vectors were constructed to evaluate the efficacy of

*UpU6* promoters in targeted gene editing of *U. prolifera*. The pCAMBIA-Cas9-sgRNA plasmid was kindly provided by Prof. Donald P. Weeks and Dr. Wenzhi Jiang from the University of Nebraska [54]. Following double digestion with *Mun*I and *Hin*dIII (New England Biolabs, Ipswich, MA, USA), the resulting ∼5.0 kb fragment was gel-purified and subsequently fused to the sgRNA cassette designed in this study, together with a ∼300 bp 5’ upstream region from one of the three *UpU6* copies via seamless assembly cloning.

All primers used were listed in Table S1. Each resulting plasmid was transformed into *E. coli* TOP10 competent cells, and plasmids were extracted using the TIANpure Midi Plasmid Kit (Tiangen Biotech Co., Ltd., Beijing, China) and submitted to Sangon Biotech Co., Ltd. for sequencing confirmation.

### 2.4 Genetic transformation in U. prolifera

Based on the genetic transformation protocol established in *U. prolifera* [18], a comprehensive assessment was conducted on the ability of the *UpU6* promoter to drive mRNA or snRNA transcription in transgenic *U. prolifera*, as well as its capacity to mediate gene editing. Transformation was performed using a Biolistic PDS-1000/He Particle Delivery System (Bio-Rad, Hercules, CA, USA). Before the bombardment, tubular thalli were sectioned into segments < 2 cm in length using a sterile blade. The segments were arranged on sterile filter paper with the majority concentrated in the central region to maximize bombardment efficiency. For each construct, 1.0 μg of plasmid DNA was adhered onto 0.6 μm gold particles according to the manufacturer’s protocol. Bombardment parameters were standardized across all treatments: 1100 psi rupture disk pressure and a target distance of 6 cm. Each plasmid was delivered in three independent bombardment experiments, serving as biological replicates.

For experiments detecting introduced gene transcription (mRNA or snRNA), healthy thalli undergoing vegetative growth were used. For gene editing assays, thalli subjected to several months of nutrient deprivation were employed. For each experiment, thalli from the same batch of cultures was collected, with equal amounts used for each bombardment. The distribution density of somatic cells on the algal surface was estimated under a microscope, and transformation was performed on the 3rd day after nutrient resupply to induce somatic *in situ* regeneration (SISR) [51,55]. Following bombardment, algal segments in all experiments were incubated at 16°C in darkness for 8–12 h to allow recovery before being returned to normal conditions.

### 2.5 Genetic transformation in Agrobacterium

Each of the three prokaryotic promoter-probe vectors and the promoterless construct was transformed into competent *A. tumefaciens* LBA4404 cells following the manufacturer’s protocol. Transformants were cultured at 28°C for 48 h on YEB medium supplemented with 50 μg/mL kanamycin. Positive colonies were identified by PCR using designed primers (Table S1) and subsequently subjected to histochemical staining for GUS expression detection.

### 2.6 Quantitative assay for GUS gene expression in U. prolifera

Quantitative GUS activity assays in *U. prolifera* were performed following established protocols [18]. Algal protein extracts were prepared 48 h post-transformation. For each sample, approximately 100 mg of thallus tissue was ground to a fine powder in liquid nitrogen and homogenized in 1 mL of GUS extraction buffer (50 mM Na2HPO4, 10 mM Na2EDTA, 0.1% Triton X-100, and 10 mM β-mercaptoethanol). The homogenate was centrifuged at 10,000 × g for 15 min at 4°C, and the supernatant was transferred to a fresh tube and maintained on ice for subsequent analysis. Soluble protein concentration was determined using the Bradford Protein Quantification Kit (Tiangen Biotech Co., Ltd.).

Transient GUS activity was measured by fluorimetric assay using 4-methylumbelliferyl-β-D-glucuronide (MUG) as substrate [56]. Each reaction mixture contained 100 μL of protein extract and 400 μL of pre-warmed assay buffer (50 mM Na2HPO4, 10 mM Na2EDTA, 0.1% Triton X-100, 10 mM β-mercaptoethanol, and 2 mM MUG). Reactions were incubated at 37°C, and 100 μL aliquots were withdrawn at 0, 15, 30, 45, and 60 min, immediately mixed with 900 μL of stop buffer (0.2 M Na2CO3), and stored in the dark at room temperature. Fluorescence intensity was determined using a multi-mode microplate reader (Infinite M1000 PRO, Tecan, Männedorf, Switzerland) with excitation at 365 nm and emission at 455 nm. Each sample was assayed in triplicate.

Statistical analyses were performed using GraphPad Prism 5.0 (GraphPad Software Inc., San Diego, CA, USA). Significant differences between groups were determined by one-way ANOVA followed by post-hoc t-tests, with *p* < 0.05 considered statistically significant.

### 2.7 Detection of heterologous U6 snRNA transcription in U. prolifera

Following bombardment with vectors carrying the *UpU6* promoter-*AtU6* cassette and recovery in darkness at 16°C for 8 h, algal thalli were transferred to normal conditions for 24 h. Total RNA was extracted from each sample using pBIOZOL Reagent (Bioflux, Beijing, China). RNA concentration and purity were assessed using a NanoDrop 2000 spectrophotometer (Thermo Fisher Scientific, Waltham, MA, USA). First-strand cDNA was synthesized by poly(A) tailing using the miRNA 1^st^ Strand cDNA Synthesis Kit (Vazyme, Nanjing, China). PCR was performed using cDNA as template with the *AtU6*-specific primer CCG-AtU6sn-F and the universal primer UniQ-R supplied in the kit (Table S1), using ApexHF HS DNA Polymerase FS Master. Subsequent steps, including product purification, A-tailing, cloning into pGEM-T vector, screening of positive *E. coli* clones, and sequencing, were performed as described previously.

### 2.8 Histochemical staining for GUS expression detection in Agrobacterium

For each prokaryotic promoter-probe construct and the promoterless negative control, kanamycin-resistant *Agrobacterium* colonies that tested positive by PCR were selected and cultured in YEB medium to an OD600 of 0.80. Histochemical staining for β-glucuronidase (GUS) activity was performed following a published protocol [56]. Briefly, 500 μL of bacterial culture was mixed with 200 μL of GUS staining solution containing 50 mM sodium phosphate buffer (pH 7.0), 0.1% Triton X-100, 2 mM K3[Fe(CN)6], 2 mM K4[Fe(CN)6], 10 mM EDTA, and 2 mM 5-bromo-4-chloro-3-indolyl-β-D-glucuronide (X-Gluc; Sangon Biotech Co. Ltd.). Samples were incubated in the dark at 37°C overnight, and color development was documented using a digital camera. The non-transformed *A. tumefaciens* LBA4404 served as the blank control.

### 2.9 Gene editing and detection of the UpAPT gene

To enable unambiguous assessment of editing events, a haploid gametophyte previously derived from the *U. prolifera* S096 strain [51] was employed for gene cloning and editing in this study. Using primers UpAPT-F/R (Table S1), the *APT* gene of *U. prolifera* (*UpAPT*) encoding adenine phosphoribosyltransferase was amplified from genomic DNA with ApexHF HS DNA Polymerase FS Master Mix. The cycling conditions consisted of an initial denaturation at 94°C for 30 s, followed by 35 cycles of 98°C for 10 s, 53°C for 15 s, and 72°C for 15 s. Based on the resolved *UpAPT* gene structure [20], sgRNA targets for *UpAPT* gene editing were designed using CRISPR-P 2.0 [57]. To evaluate the cleavage activity of sgRNAs, a partial *UpAPT* sequence containing the target sites was amplified using primers dUpAPT-F/R (Table S1). sgRNAs were transcribed *in vitro* using the sgRNA In Vitro One-step Transcription Kit (Novoprotein Scientific Inc.). The cleavage activity of sgRNAs was validated *in vitro* using the sgRNA Screening and Genotype Confirmation Kit (Novoprotein Scientific Inc.). Following the cleavage reaction, 1.5% agarose gel electrophoresis was performed to assess Cas9-mediated sgRNA cleavage efficiency. Valid sgRNAs were selected and employed, as described above, to construct *UpAPT* gene editing vectors by fusing the *UpU6* promoter.

Following Biolistic bombardment, the samples were initially recovered in the dark at 16°C for 8 h, and then transferred to normal culture conditions. Ten days later, when most regenerated seedlings had developed to the 12–16-cell stage, the proportion of cells undergoing SISR was assessed by microscopy, and the seedlings were selected with 2-fluoroadenine (2-FA) at a final concentration of 50 μM. Selective pressure was maintained for 4–6 weeks, with the VSE medium replaced every 5–7 days. All surviving seedlings were individually selected, and genomic DNA was extracted for PCR amplification with primers eUpAPT-F/R (Table S1) to detect *UpAPT* mutations. For samples that failed to produce the expected amplicons, a combined strategy of tiling PCR and long-range PCR was implemented. Q5 High-Fidelity DNA Polymerase (New England Biolabs, USA) together with the primer pairs XUpAPT-F/R, YUpAPT-F/R, and maxAPT-F/R (Table S1) were employed to screen for potential deletions or large insertions.

## 3. Results

### 3.1 Cloning and alignment of UpU6 genes with 5’ upstream sequences

A BLASTN search was performed using the complete 101 bp *CrU6* sequence from *C. reinhardtii* (GenBank: X71486.1) against three available *U. prolifera* genome assemblies. Within the chromosome-level assembly from a floating YSGT sample (GenBank: JALAAN000000000.1), three identical copies of the *UpU6* gene, each approximately 105 bp in length, were identified and designated *UpU6-1*, *UpU6-2*, and *UpU6-3*. *UpU6-1* was mapped to chromosome 3 (GenBank: JALAAN010000003.1), whereas *UpU6-2* and *UpU6-3* were both located on chromosome 1 (GenBank: JALAAN010000001.1). These three gene copies were also detected in a second YSGT sample dataset (GenBank: SDUY00000000.1). In contrast, only two *UpU6* copies were identified in the genome assembly from an attached sample collected in Japan (GenBank: BRCE00000000.1), designated *UpU6-4* (from BRCE01000010.1) and *UpU6-5* (from BRCE01000005.1). All five gene copies, *UpU6-1* to *UpU6-5*, shared identical sequences.

To assess the copy number of *UpU6*, a maximum likelihood (ML) phylogenetic tree was constructed using sequences encompassing each *UpU6* copy and approximately 2.0 kb of upstream flanking sequence. As shown in Fig. S1, all sequences clustered into three distinct clades, indicating that only three *UpU6* copies exist and that *UpU6-4* and *UpU6-5* are homologous to *UpU6-2* and *UpU6-1*, respectively. This might be due to the incomplete genomic resources of the attached samples, resulting in the failure to identify the homolog of *UpU6-3*.

**Fig. S1.**
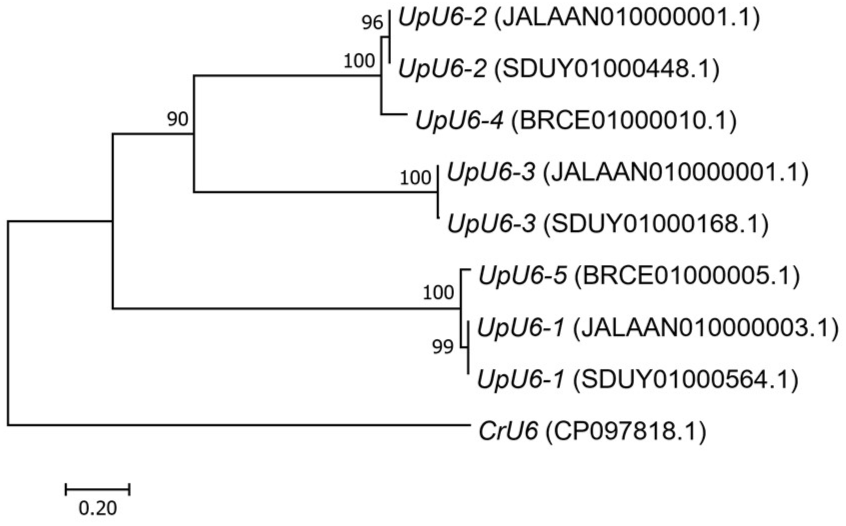
ML phylogenetic tree based on nucleotide sequences of each *UpU6* copy and approximately 2.0 kb of upstream flanking sequence. GenBank accession numbers indicate the specific contig, scaffold, or chromosome containing each sequence. Bootstrap support values are shown at the nodes.

Based on the available genome assemblies, primers were designed to amplify each *UpU6* copy together with 1.0–1.4 kb of its 5’ upstream sequence. Amplification yielded fragments of 1448 bp (GenBank: PV637427.1), 1499 bp (GenBank: PV637428.1), and 1644 bp (GenBank: PV637429.1) using primer pairs UpU6-1-F/R, UpU6-2-F/R, and UpU6-3-F/R, respectively. Given that the nature of S096 strain utilized in this study has been identified as the floating ecotype, attempts to amplify *UpU6-4* and *UpU6-5* using the corresponding primer pairs yielded no PCR products, as expected. Sequence polymorphisms between each amplified fragment and its corresponding reference sequence (JALAAN000000000.1) were assessed. As shown in Fig. S2, the 1448 bp and 1499 bp fragments were completely identical to their respective references, whereas the 1644 bp fragment contained two deletions and several single nucleotide polymorphisms (SNPs) relative to the reference. These results confirm that all three *UpU6* copies (*UpU6-1* to *UpU6-3*), along with their upstream regions, were successfully amplified from the YSGT floating sample.

**Fig. S2.**
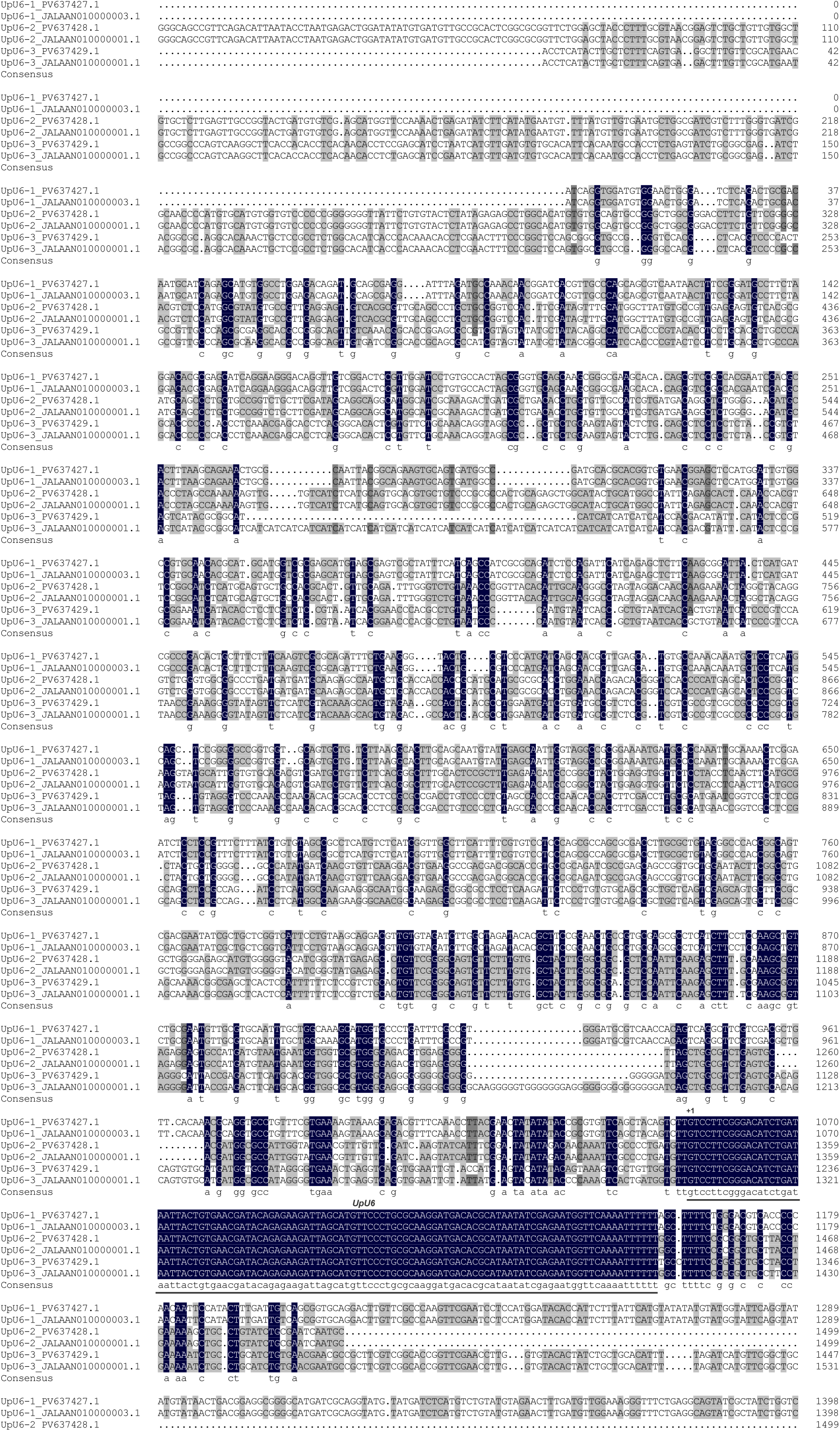
Alignment of amplified *UpU6* copies (*UpU6*-1 to *UpU6-3*) with their corresponding reference sequences from assembly JALAAN000000000.1. Each sequence encompasses the *U6* snRNA gene and its 5’ upstream flanking region. The black bar denotes the *UpU6* gene. The transcription start site (TSS) is designated as +1.

To delineate the *UpU6* genes and identify conserved regulatory elements in plants, sequences of the three copies were aligned with orthologs from representative species, including the chlorophyte *C. reinhardtii*, the liverwort *Marchantia polymorpha*, and multiple species of angiosperms. The *U6* snRNA gene sequences exhibited a high degree of conservation across species, with the *UpU6* gene being approximately 105 bp in length and showing complete sequence identity among the three copies. Each *UpU6* gene initiated with a guanine (G) residue and terminated with a thymine (T)-rich tract, consistent with the conserved architecture of *U6* snRNA genes and potentially serving as a transcription termination signal. In contrast, the upstream regions showed considerable interspecific variation. Notably, a TATA box was identified approximately 30 bp upstream of each *U6* snRNA gene, suggesting its function as a core promoter element. However, the upstream sequence element (USE, consensus RTCCCACATCG), which is typically positioned around −60 and recognized as a crucial regulatory element for U6 snRNA transcription in seed plants [28,58], appeared to be absent in liverwort and chlorophyte (Fig. 1).

**Fig. 1.**
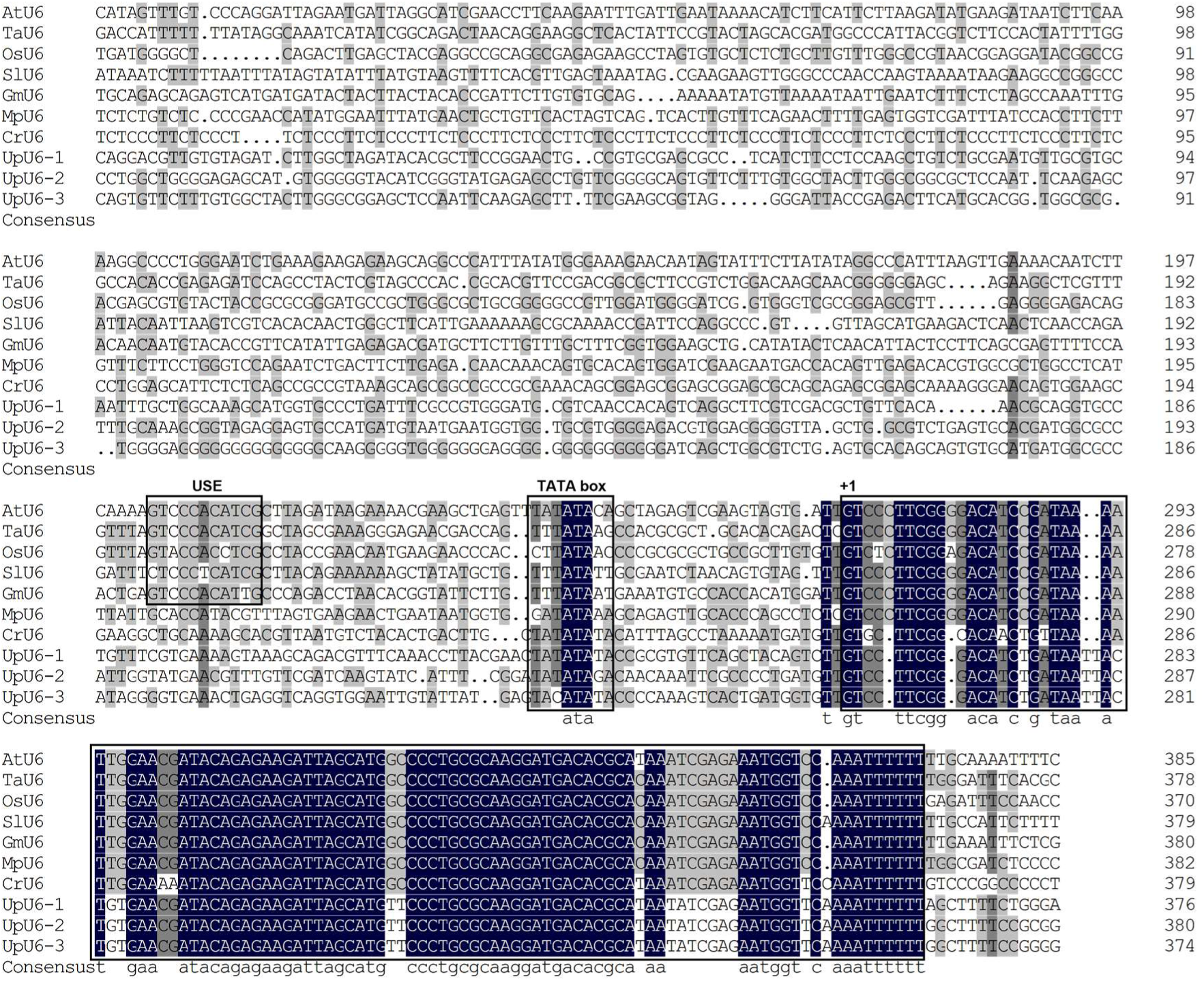
Multiple sequence alignment of *U6* snRNA genes and their upstream regions from *U. prolifera* and representative species. Conserved regulatory elements—USE, TATA box, and *U6* snRNA genes sequences—are highlighted with black boxes, respectively. The TSS is designated as +1. Representative species and GenBank accession numbers are as follows: AtU6 (*Arabidopsis thaliana*, X52528.1); TaU6 (*Triticum aestivum*, CM031188.1); OsU6 (*Oryza sativa*, AP014967.1); SlU6 (*Solanum lycopersicum*, X51447.1); GmU6 (*Glycine max*, NC038255.2); MpU6 (*Marchantia polymorpha*, AB902572.1); CrU6 (*Chlamydomonas reinhardtii*, X71486.1).

### 3.2 Cis-regulatory element analysis of 5’ upstream sequences of UpU6 genes

Using PlantCARE, potential *cis*-regulatory elements within the 5’ upstream regions of each *UpU6* copy were predicted. The results showed that all copies contained the core promoter element TATA box, along with various predicted *cis*-regulatory elements associated with light, hormone, or stress responses. Notably, the three copies exhibited distinct differences in sequence composition and element distribution. Methyl jasmonate (MeJA)-responsive elements, including the TGACG-motif and CGTCA-motif, were abundant in the 5’ upstream regions of *UpU6-1* and *UpU6-3*, but completely absent from *UpU6-2*. Conversely, *UpU6-2* contained a substantially higher number of abscisic acid (ABRE)-and light (G-box)-responsive elements compared with the other two copies (Figs. S3–S5). However, the functional contributions of individual motifs remain to be experimentally validated. These results raise the possibility that transcription of individual *UpU6* copies may be differentially regulated by specific environmental cues.

**Fig. S3.**
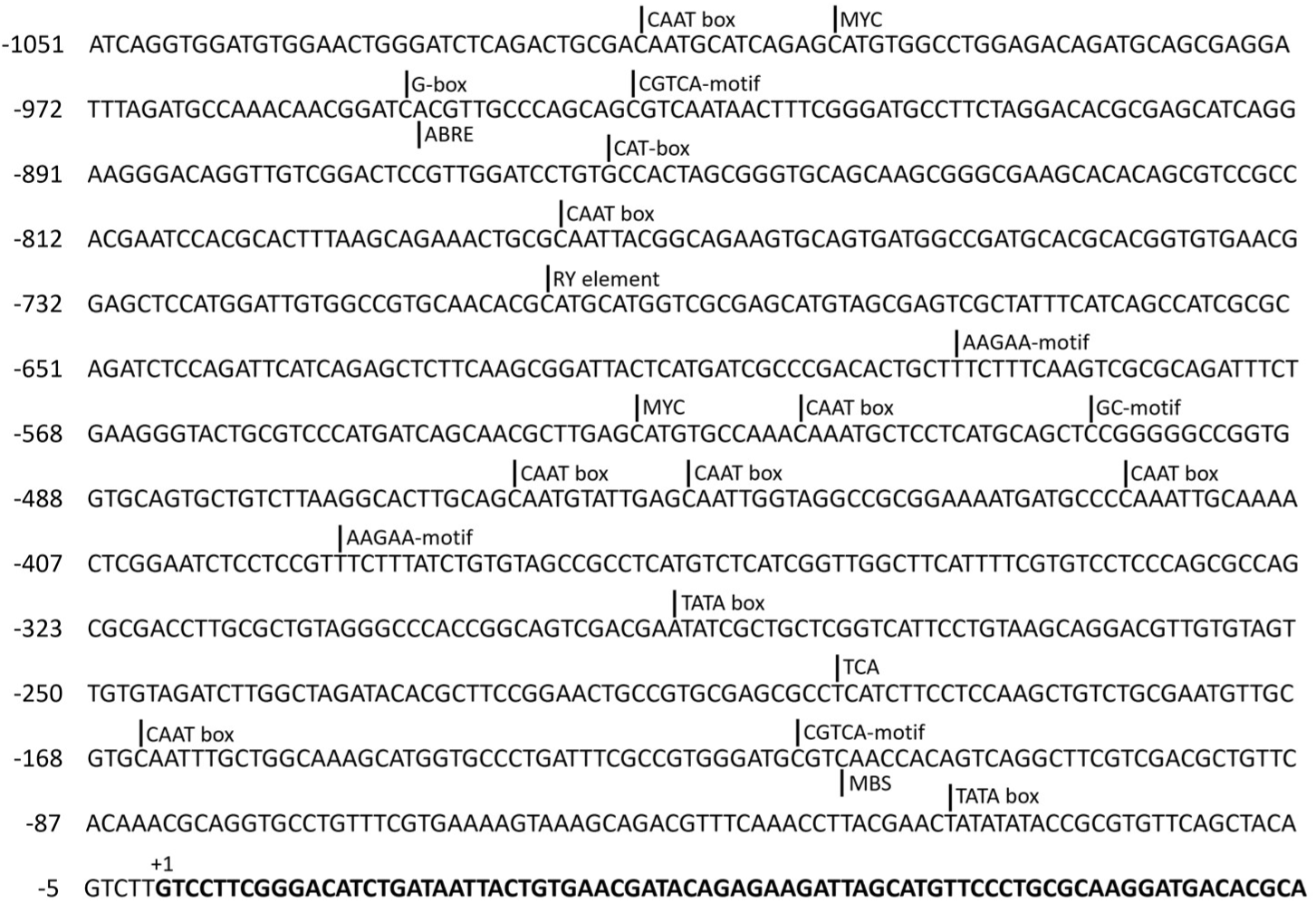
Predicted *cis*-regulatory elements in the 5’ upstream region of *UpU6-1*. Their positions relative to the TSS of *UpU6* (+1) are indicated. Vertical lines denote the first base of each element.

**Fig. S4.**
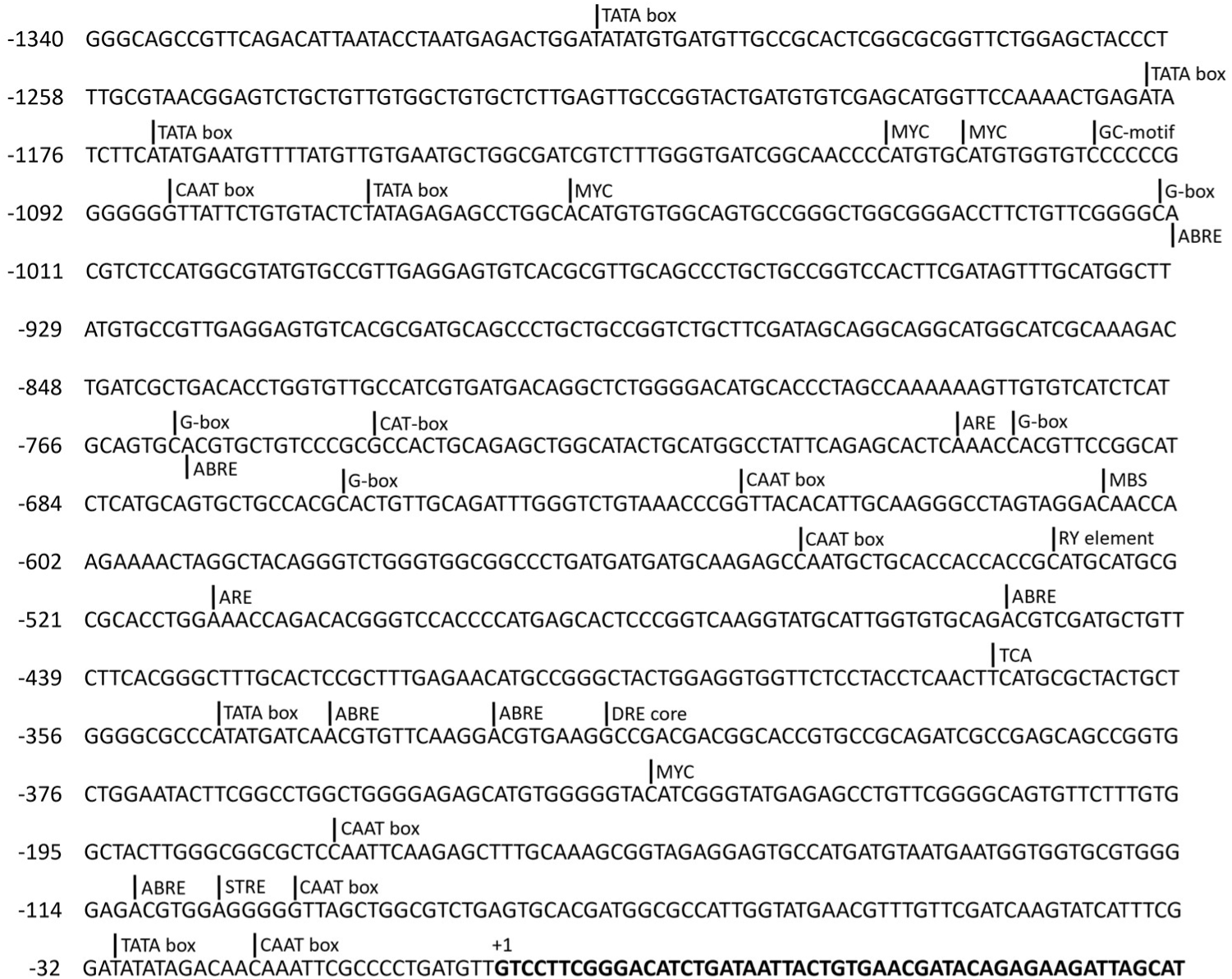
Predicted *cis*-regulatory elements in the 5’ upstream region of *UpU6-2*. Their positions relative to the TSS of *UpU6* (+1) are indicated. Vertical lines denote the first base of each element.

**Fig. S5.**
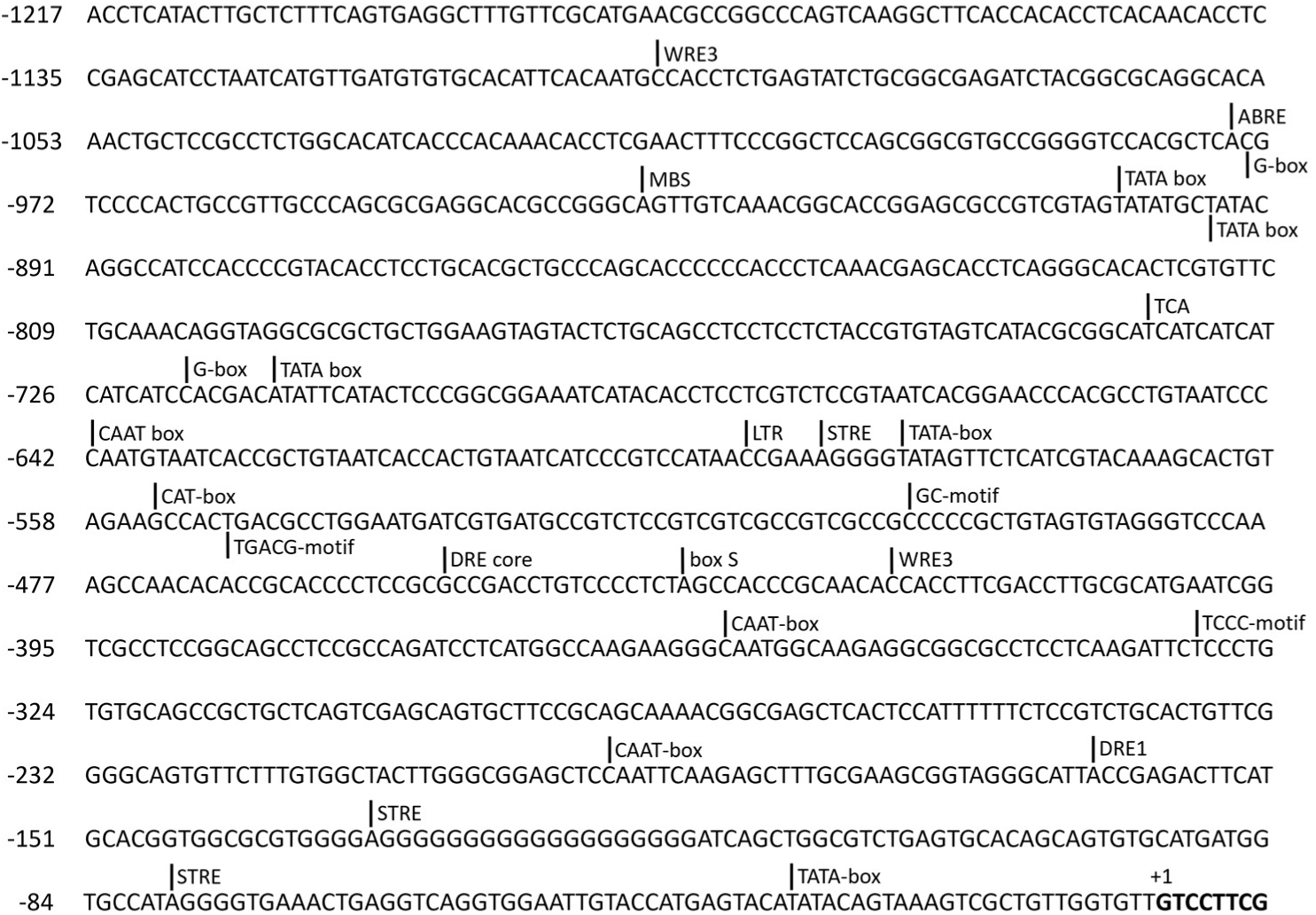
Predicted *cis*-regulatory elements in the 5’ upstream region of *UpU6-3*. Their positions relative to the TSS of *UpU6* (+1) are indicated. Vertical lines denote the first base of each element.

### 3.3 Detection of mRNA transcription driven by UpU6 promoters in U. prolifera

The activity of *UpU6* promoters in driving mRNA transcription was examined in *U. prolifera*, a multicellular eukaryotic seaweed. Using pBI221 as the backbone vector, three eukaryotic promoter-probe vectors—p221U6-1P-GUS, p221U6-2P-GUS, and p221U6-3P-GUS—were constructed by replacing the native CaMV35S promoter with ∼1.0 kb of the 5’ upstream region from each *UpU6* copy (Fig. 2a).

**Fig. 2.**
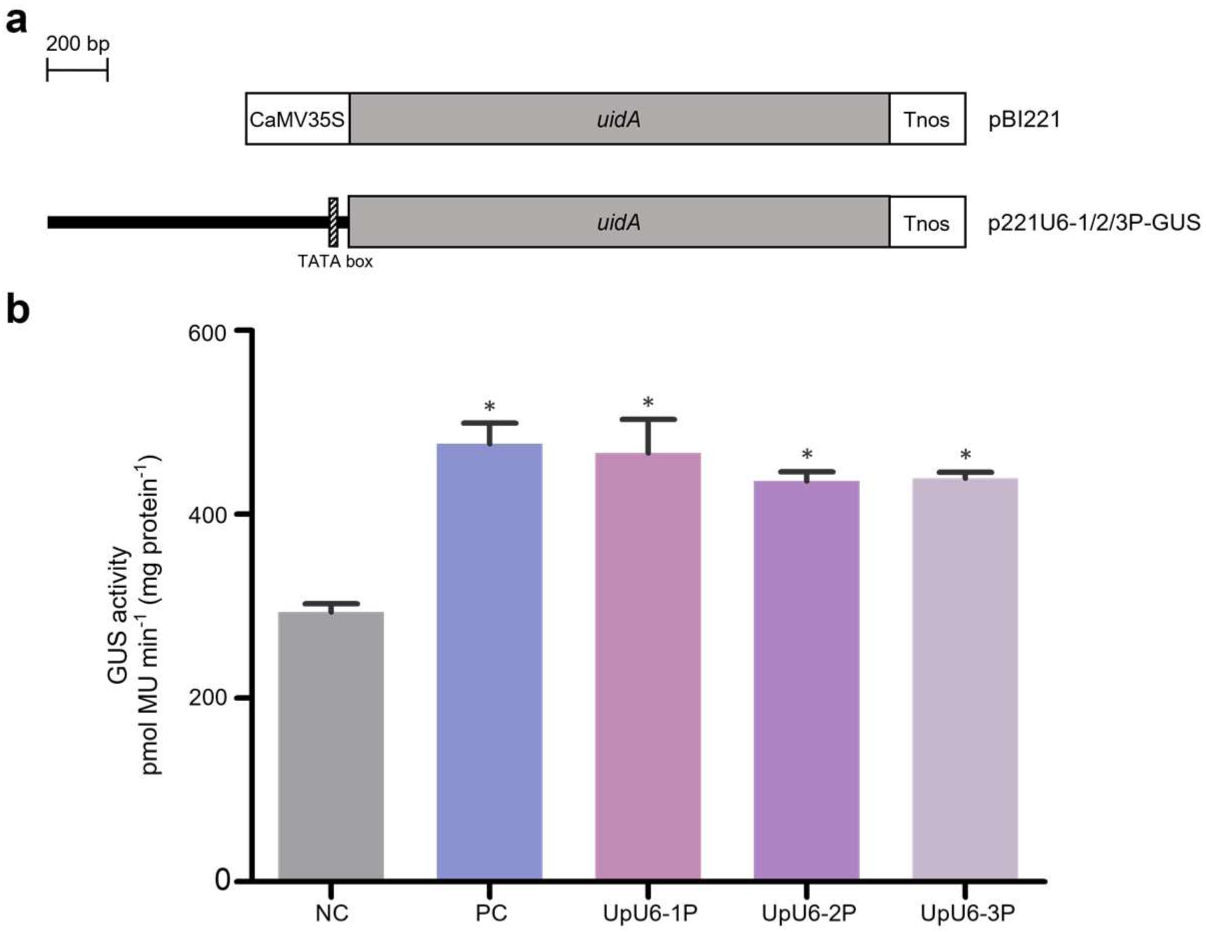
Functional validation of *UpU6* promoters in driving mRNA transcription in *U. prolifera*. (a) Schematic diagram of vectors used for mRNA transcription driven by *UpU6* promoters in *U. prolifera*. The thick black bar represents *UpU6* promoters. CaMV35S, *uidA*, and *Tnos* represent the CaMV35S promoter, *GUS* reporter gene, and *nos* terminator, respectively. (b) The expression levels of *GUS* driven by different promoters. Error bars indicate standard deviation (SD) from three independent biological replicates. Asterisks indicate significant differences (\**p* < 0.05). NC represents the negative control. PC represents the positive control. UpU6-1P/2P/3P represent the experimental groups using the *UpU6-1/2/3* promoters with 1.0 kb truncation.

The particle bombardment method was employed to introduce the above-mentioned reporter vectors into *U. prolifera*, with pBI221 as the positive control and untransformed algae as the negative control. At 72 h post-transformation, GUS activity was quantitatively assessed across all groups. As shown in Fig. 2b, both the positive control and all *UpU6* promoter treatment groups differed significantly from the negative control (*p* < 0.05), indicating that all three *UpU6* promoters can drive mRNA transcription in *U. prolifera*. However, GUS activity levels were comparable among the three *UpU6* promoters, with no significant difference relative to the positive control.

### 3.4 Detection of heterologous U6 snRNA transcription in U. prolifera

To validate the ability of the *UpU6* promoters to drive small RNA transcription in *U. prolifera*, a 104-bp copy of the *AtU6* gene was amplified from the *A. thaliana* Columbia ecotype. This fragment was then separately fused in tandem with three 300-bp truncated *UpU6* promoter fragments to construct vectors p221UpU6-1At, p221UpU6-2At, and p221UpU6-3At (Fig. 3a). After each vector was introduced into *U. prolifera* thalli via Biolistic, total RNA was extracted at 24 h post-transformation, and small RNAs were reverse-transcribed. The cDNA was subsequently subjected to PCR with the designed *AtU6*-specific primer CCG-AtU6sn-F (Fig. 3a) and a kit-provided universal primer. The PCR products were cloned and sequenced, and representative sequences were selected for alignment (Fig. 3b).

**Fig. 3.**
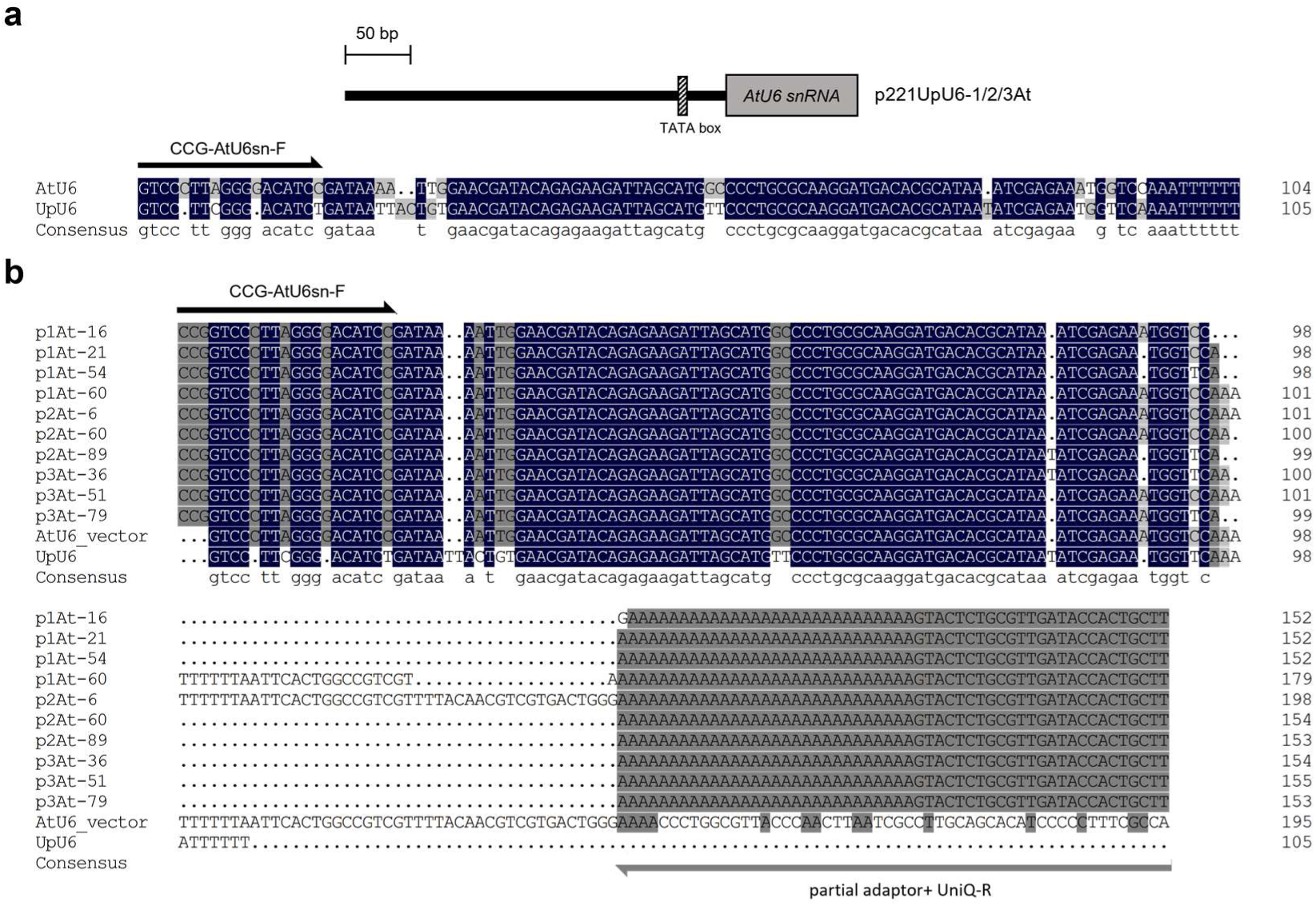
Functional validation of *UpU6* promoters in driving heterologous U6 snRNA transcription in *U. prolifera*. (a) Schematic diagram of vectors used for *in vivo* transcription of snRNA driven by *UpU6* promoters in *U. prolifera* and the binding region of the *AtU6*-specific 5’ primer CCG-AtU6sn-F. Black thick bar represents the promoter. Black arrow indicates the primer binding site and the extension direction. (b) Sequence alignment of the representative amplified transcripts with *AtU6* and *UpU6*. AtU6_vector denotes the sequence of *AtU6* and its 3’ flanking vector regions within the constructed p221UpU6-1/2/3At vectors. Partial adaptor+UniQ-R represents a region that includes the tailing sequence added by the reverse transcription kit and the universal primer UniQ-R. p1/2/3At represent the clones derived from the experimental groups using the *UpU6-1/2*/*3* promoters, respectively.

The results showed that, first, for each *UpU6* promoter, at least one clone accurately reflected the target *AtU6* sequence (p1At-60, p2At-6, p2At-60, and p3At-51), indicating that all three *UpU6* promoters are capable of driving small RNA transcription. It should be noted that the majority of the 5’ ends of all the clones fully exhibit the characteristics of *AtU6*, while the 3’ ends of some clones (p1At-21, p1At-54, p2At-89, p3At-36, p3At-79) were homologous to *UpU6*. This was very likely to be the result of chimeric PCR, as the 3’ primer was a universal primer rather than the *AtU6*-specific primer. Second, most transcripts terminated at the poly(T) tract, a hallmark of Pol III-dependent termination, suggesting that the *UpU6* promoter is likely to drive the snRNA transcription by recruiting Pol III rather than Pol II. Very few clones (p1At-60 and p2At-6) exhibited transcriptional read-through, where several dozen base pairs of vector sequence immediately downstream of *AtU6* were also transcribed, suggesting that the original poly(T)-downstream sequence—replaced by vector sequence—may contain additional auxiliary termination signals.

### 3.5 Detection of mRNA transcription driven by UpU6 promoters in Agrobacterium

To determine whether the three *UpU6* promoters can drive mRNA transcription in prokaryotic cells, approximately 300 bp 5′ upstream regions of each *UpU6* copy were amplified. Using seamless assembly cloning, three prokaryotic promoter-probe vectors were constructed by fusing each *UpU6* promoter fragment to the *GUS* reporter gene in a modified pCAMBIA1301 backbone: p1301U6-1P-GUS, p1301U6-2P-GUS, and p1301U6-3P-GUS (Fig. 4a).

**Fig. 4.**
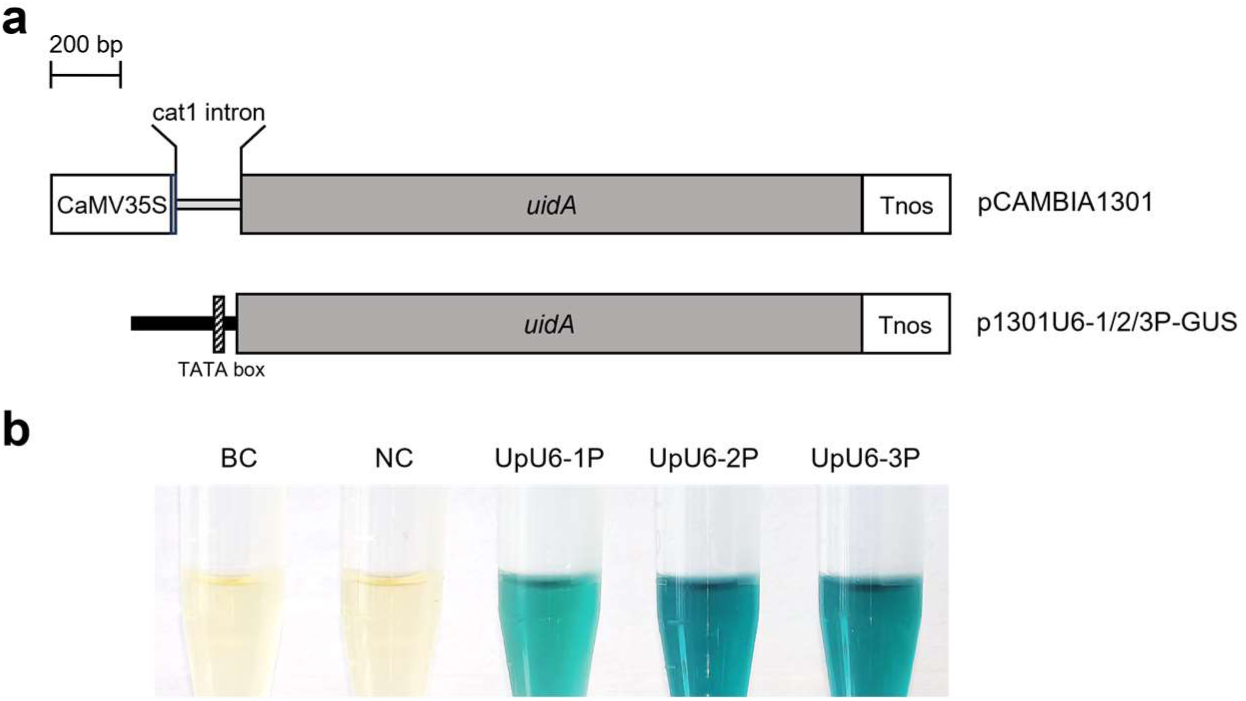
Functional validation of *UpU6* promoters in supporting GUS reporter expression in *Agrobacterium*. (a) Schematic diagram of vectors used for mRNA transcription driven by *UpU6* promoters in prokaryotes. Thick black bars represent *UpU6* promoters. CaMV35S, *uidA*, and *Tnos* represent the CaMV35S promoter, *GUS* reporter gene, and *nos* terminator, respectively. The cat1 intron indicates the intron originally present within the *GUS* gene in the pCAMBIA1301 vector. (b) The staining of GUS driven by *UpU6* promoters in *Agrobacterium*. BC represents the blank control. NC represents the negative control. UpU6-1P/2P/3P represent the experimental groups using the UpU6-1/2/3 promoters with 300 bp truncation.

Each constructed plasmid was individually transformed into *A. tumefaciens* LBA4404. Kanamycin-resistant transformants were verified by PCR, and bacterial cultures were subsequently subjected to histochemical staining to detect GUS expression. Blue staining was observed in all three treatment groups carrying the respective *UpU6* promoter regions, whereas no blue staining was detected in either the promoterless control or the non-transformed *A. tumefaciens* LBA4404 control (Fig. 4b). These results indicated that all three *UpU6* promoter regions were capable of supporting GUS reporter expression in prokaryotic cells.

### 3.6 Gene editing and molecular identification of the UpAPT gene

To enable precise gene editing, the 1387-bp genomic DNA sequence of the *UpAPT* gene—from start codon to stop codon—was determined from the haploid male gametophyte genome of strain S096 (GenBank: PZ397805.1), and SNPs were identified by comparing with known *UpAPT* sequences (Fig. S6). Based on the reported structure of the *UpAPT* gene, six candidate targets of sgRNA (sg1 to sg6) were designed within the third exon (Fig. 5a). Following PCR amplification, the corresponding sgRNAs were synthesized by *in vitro* transcription. *In vitro* cleavage assays (Fig. 5b) showed that all six sgRNAs efficiently guided Cas9-mediated cleavage at their respective target sites, with comparable efficiencies among the different guides. Three of these (sg1, sg2, and sg3) were selected to construct vectors for subsequent gene editing experiments.

**Fig. S6.**
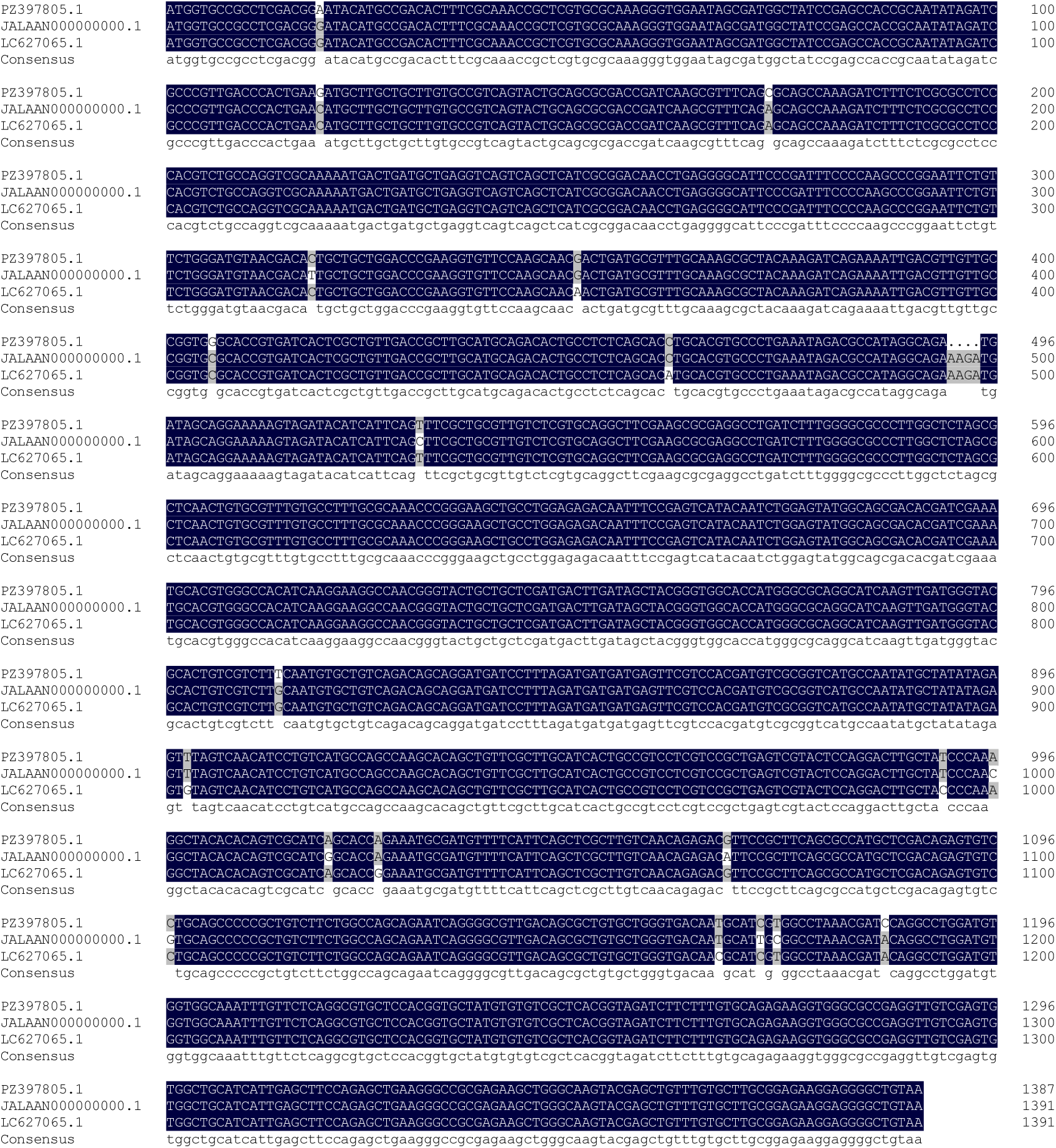
Alignment of *UpAPT* genomic sequence obtained in this study with reference sequences. PZ397805 was generated in this study and deposited in GenBank. JALAAN000000000.1 and LC627065.1 were reference sequences from GenBank.

**Fig. 5.**
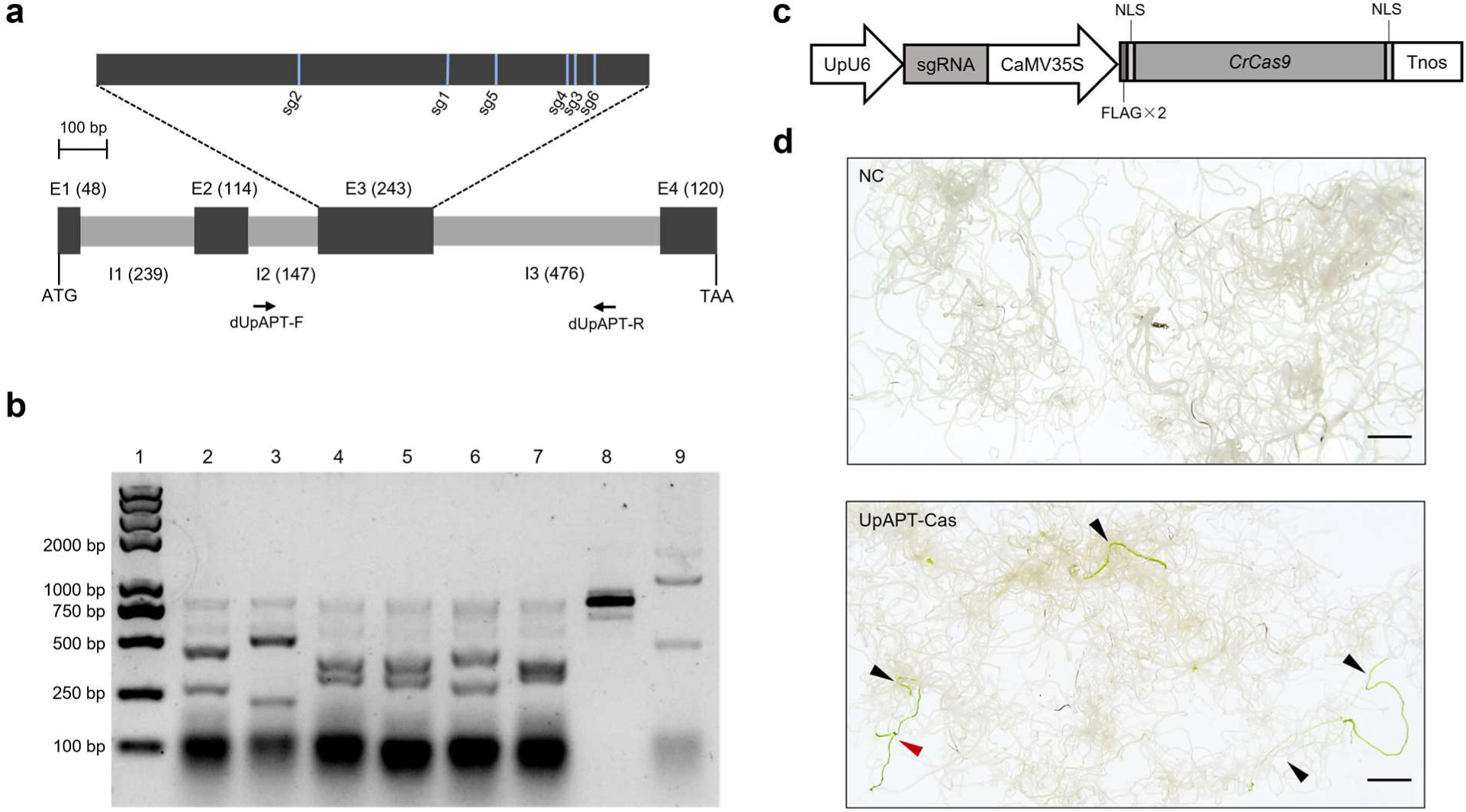
CRISPR-Cas9-mediated editing of *UpAPT* in *U. prolifera*. (a) Schematic diagram of the *UpAPT* gene structure and the locations of designed sgRNA target sites. Black boxes (E1–E4) represent exons, gray boxes (I1–I3) represent introns, with the length of each region (in bp) indicated in parentheses. (b) *In vitro* cleavage assay of designed sgRNAs targeting *UpAPT*. Lane 1 represents DNA marker. Lanes 2–7 represent substrate DNA incubated with Cas9 and the corresponding sgRNAs (sg1–sg6), respectively. Lane 8 represents substrate DNA only. Lane 9 represents the positive control provided in the kit. (c) Schematic diagram of CRISPR-Cas9 vectors for gene editing of *UpAPT* in *U. prolifera.* NLS represents the nuclear localization signal. FLAG represents the FLAG epitope tag. (d) Survival of negative control and CRISPR-Cas9 transformants under 2-FA selection. The negative control (NC) refers to thalli transformed with bare gold particles; UpAPT-Cas9 refers to thalli transformed with the CRISPR-Cas9 constructs targeting *UpAPT*. Black arrows indicate separated surviving individuals. Red arrow indicates multiple individuals occurring in clusters. Scale bars = 0.5 cm.

Using the Seamless Assembly Cloning Kit, five CRISPR-Cas9 vectors were constructed by combining different *UpU6* promoter copies (*UpU6-1* to *UpU6-3*) with sgRNAs (sg1 to sg3) (Fig. 5c), designated as pU6-1-sg1, pU6-1-sg2, pU6-1-sg3, pU6-2-sg3, and pU6-3-sg3, respectively. On day 3 after inducing SISR in the male gametophytes of *U. prolifera* strain S096, the vectors were introduced by particle bombardment, with a negative control group bombarded with naked gold particles. After 10 days of cultivation, most regenerated seedlings (> 70%) in both the treatment and negative control groups had developed to the 12–16-cell stage. At this point, 2-FA was added at a final concentration of 50 μM for selection. After two weeks, the algal thalli in all groups gradually died and turned white to the naked eye. Following an additional 2–4 weeks of selection, green thalli were observed in all experimental groups, totaling 52 individuals across all groups, with most appearing separately and few occurring in clusters, whereas no viable algae were observed in the negative control group (Fig. 5d).

To identify mutations in the target gene in all 2-FA-resistant individuals, we initially examined the target regions and subsequently expanded PCR amplification to the broader *UpAPT* genomic region because the observed mutation patterns were more complex than initially anticipated. Amplicons were directly obtained from 32 samples, while 16 samples required extra tiling PCR or long-fragment PCR. The remaining four samples always failed to amplify, suggesting possible large-scale structural variations (SVs). Mutations were detected within the exon regions of the *UpAPT* gene in all 48 sequenced samples (Fig. 6), demonstrating that all three *UpU6* promoters effectively drove sgRNA transcription and guided

**Fig. 6.**
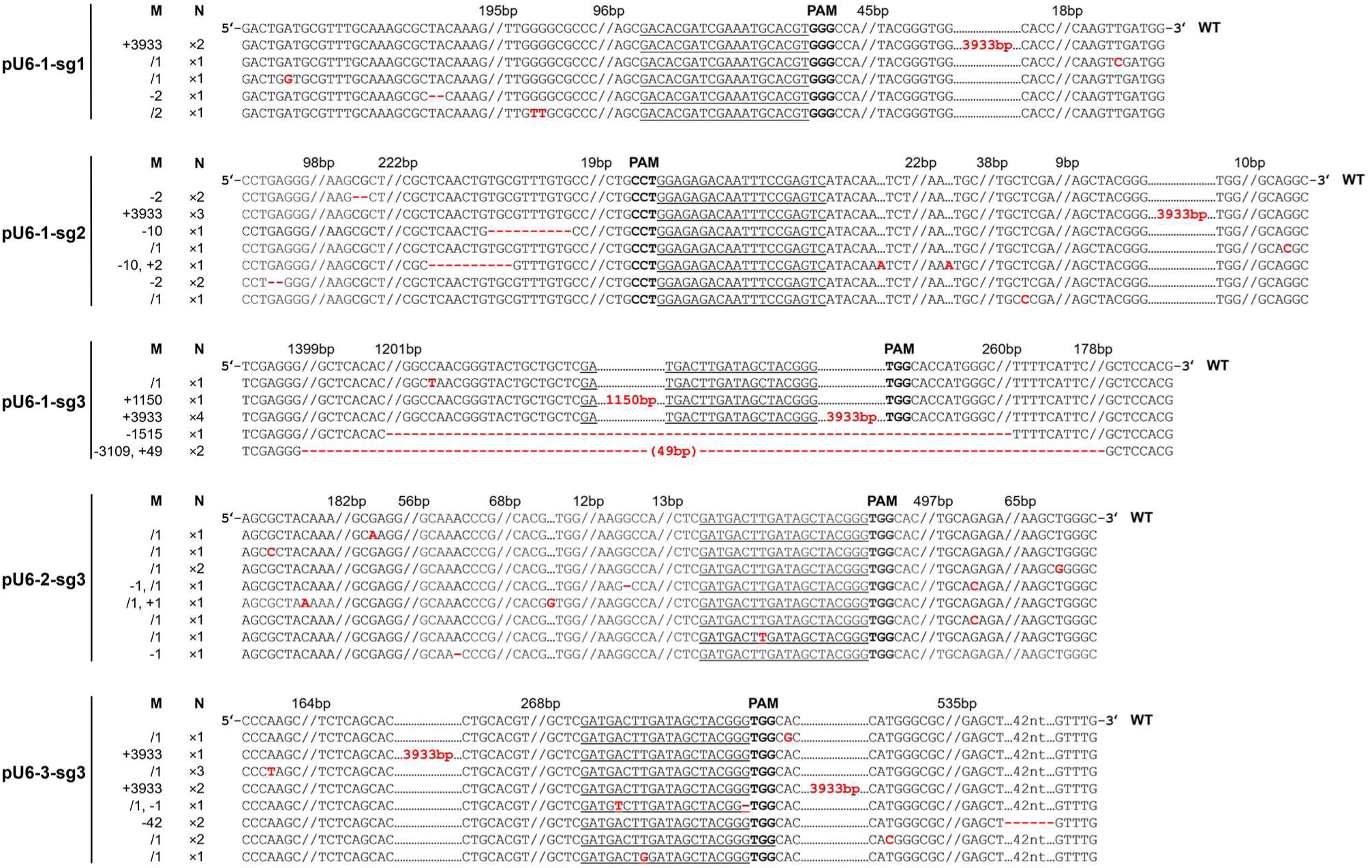
Characterization and distribution of CRISPR-Cas9-induced mutations at the *UpAPT* locus. WT represents the wild-type. The underlined sequences indicate sgRNA target sequences, while bold fonts represent protospacer adjacent motif (PAM) sequences. Red annotations indicate gene-editing mutations and associated sequence changes. Red nucleotides represent inserted or substituted bases, whereas red hyphens indicate nucleotide deletions. Red numbers (bp) indicate the length of inserted sequences, and bracketed red numbers (bp) denote the length of inserted sequences of unknown origin within large deletion regions. Double slashes indicate omitted sequences, with the numbers above representing the length of omitted sequences (in bp). Column M indicates the mutation type and the number of nucleotides involved, where “–” denotes deletion, “+” denotes insertion, and “/” denotes substitution. Column N represents the number of mutant individuals sharing the same edited genotype.

Cas9-mediated editing of the *UpAPT* gene. The editing efficiency was evaluated using two methods: the mutation-positive fraction among resistant thalli was 92.3% (48/52); and the editing frequency calculated from estimated starting cell number ranged from 1.88 × 10⁻⁶ to 4.07 × 10⁻⁶, with an average of 3.26 × 10⁻⁶ (Table 1).

**Table 1.** Editing events and editing efficiency of the *UpAPT* gene.

| Experimental group | 2-FA-resistant thalli | Editing events |  |  |  | Editing efficiency |  |  |
| --- | --- | --- | --- | --- | --- | --- | --- | --- |
| | | Substitution | Indels | Large insertion | Large deletion | N.D.<sup>a</sup> | By thallus<sup>b</sup> | By cell ($\times 10^{-6}$)<sup>c</sup> |
| pU6-1-sg1 | 6 | 3 | 1 | 2 | 0 | 0 | 100% (6/6) | 1.88 |
| pU6-1-sg2 | 13 | 2 | 6 | 3 | 0 | 2 | 84.6% (11/13) | 4.07 |
| pU6-1-sg3 | 10 | 1 | 0 | 5 | 3 | 1 | 90% (9/10) | 3.13 |
| pU6-2-sg3 | 10 | 8<sup>d</sup> | 3<sup>d</sup> | 0 | 0 | 1 | 90% (9/10) | 3.13 |
| pU6-3-sg3 | 13 | 8<sup>e</sup> | 3<sup>e</sup> | 3 | 0 | 0 | 100% (13/13) | 4.07 |
| In total | 52 | 21 | 13 | 13 | 3 | 4 | 92.3% (48/52) | 3.26<sup>f</sup> |

<sup>a</sup> N.D. indicates that the mutation type was not determined due to failed PCR amplification, possibly caused by large genomic rearrangements at the target locus.
<sup>b</sup> The mutation-positive fraction among resistant thalli.
<sup>c</sup> The editing frequency calculated from the starting number of surface cells.
<sup>d</sup> Two indel/substitution complex editing events were included.
<sup>e</sup> One indel/substitution complex editing event was included.
<sup>f</sup> The average values of five experimental groups.

Sequence analysis revealed that small-scale mutations, including substitutions and indels, constituted the predominant type of editing events. Notably, editing mediated by the *UpU6-2* promoter exclusively generated small-scale mutations, with no large-scale inversions or deletions detected. In contrast, large insertions of 3933 bp were identified in editing events driven by both the *UpU6-1* and *UpU6-3* promoters. Further analysis revealed that the inserted 3933-bp fragment originated from a Mutator-like element (MULE), a class of DNA transposons in the *U. prolifera* genome, and exhibited typical MULE features, including ∼111-bp Terminal Inverted Repeats (TIRs) at both ends and an 8-bp Target Site Duplication (TSD) at the insertion site. Additionally, a 1150-bp large insertion was detected in the *UpU6-1*-driven editing, which represented a truncated fragment of the 3933-bp insertion and lacked both TIRs and TSDs. Furthermore, two large deletions of 1515 bp and 3109 bp were identified in the editing results driven by the *UpU6-1* promoter. Sequence analysis revealed that the deleted regions spanned exons 1–3 of the *UpAPT* gene and part of the 5’ upstream sequence, resulting in complete loss of gene function. It should be noted that some samples grew into a cluster (Fig. 5d) and exhibited identical edited sequences. This suggested that the edited cells underwent cell division prior to SISR, thereby producing new individuals carrying the same editing event. To avoid overestimation from clonal individuals bearing identical edited genotypes, only independent editing events were counted in calculating the MULE insertion frequency. The pU6-1-sg1, pU6-1-sg2, pU6-1-sg3, pU6-2-sg3, and pU6-3-sg3 groups yielded 5, 7, 5, 8, and 8 independent editing events, respectively, corresponding to an overall MULE insertion frequency of 18.2% (6/33).

## 4. Discussion

In this study, three *U6* promoters of *U. prolifera* were identified and cloned. The functional validation results showed that in addition to driving the transcription of small RNA, these promoters could also drive protein-coding gene expression in *U. prolifera* and *Agrobacterium*. Notably, all three *UpU6* promoters can be applied to the vector-based CRISPR-Cas9 system, enabling targeted editing of the *UpAPT* gene and obtaining live mutants. These findings expand the understanding of the functional flexibility of *U6* promoters, while also enriching the toolkit of regulatory elements for the CRISPR-Cas9 genome editing platform in *U. prolifera*.

### 4.1 Conservation and evolutionary dynamics of U6 snRNA genes and promoters

U6 snRNA is a core component of the spliceosome in eukaryotes, whose sequence and structure are highly conserved [37,59]. It was found that three *UpU6* copies identified in this study shared identical sequences and retained the characteristic features of a “G” nucleotide at the 5’ end and a poly(T) tract at the 3’ end [60,61]. Although their precise lengths have not yet been determined, they fell within the range of approximately 100–120 nt for eukaryotic U6 snRNAs [62], reflecting the stringent functional constraints imposed on U6 snRNA within the catalytic core of the spliceosome [59,62]. In contrast, the copy number of *U6* snRNA genes varies considerably among eukaryotes: it was present as a single copy in the yeast *Saccharomyces cerevisiae* [59], 3–5 copies in some green algae including *C. reinhardtii* (data not shown) and *Ulva* (Blomme et al., 2025), generally a dozen or so copies in seed plants [40,63,64], and up to several hundred copies in mammals [65]. It was indicated that the *U6* snRNA gene copy number was likely associated with transcriptional demands, which were in turn strongly affected by the genome complexity and the degree of specific expression of genes [37].

U6 snRNA is typically transcribed by Pol III via the type 3 Pol III promoter [27,28]. Transcriptional regulation of this promoter type relies exclusively on upstream *cis*-acting elements, with no dependence on intragenic sequences [66]. In this study, all three *UpU6* promoters were found to contain only a conserved TATA box element located approximately -30 bp upstream of the TSS, which is consistent with the promoter architecture observed in the unicellular green alga and liverwort (Fig. 1). In contrast, the *U6* promoter of higher plants has evolved more conserved regulatory elements. For instance, in dicotyledonous plants, the *U6* promoter generally contains positionally conserved USE elements, while in monocotyledonous plants, additional monocot-specific promoter (MSP) elements with similar enhancer functions have been added [58]. This suggested that the *U6* promoter might have gradually evolved from a relatively simple core structure to a complex system containing multiple *cis*-regulatory elements during the evolution of the green plant lineage, in order to meet the more precise transcriptional regulation requirements. Furthermore, despite the identical sequences of the three *UpU6* copies, their upstream regulatory regions differed substantially in both the types and arrangements of predicted *cis*-regulatory elements, suggesting that they may be subject to distinct transcriptional regulation. However, this study only compared their activities during normal vegetative growth or SISR process, differences under specialized physiological conditions or other life history stages remain to be investigated.

### 4.2 Dual transcriptional activity and cross-system functionality of UpU6 promoters

In eukaryotes, U6 snRNA is typically transcribed by Pol III, whereas mRNA is transcribed by Pol II, generating a 7-methylguanosine (m7G) cap at the 5’ end and a 3’ poly(A) tail essential for mRNA stability and efficient translation. In this study, the *UpU6* promoters were shown to drive U6 snRNA transcription, with most products terminating at the poly(T) signal, consistent with canonical Pol III characteristics. Meanwhile, all three *UpU6* promoters exhibited ability to drive protein-coding gene expression in *U. prolifera*. These facts suggested that the *UpU6* promoters may possess dual small-RNA/mRNA transcriptional activity, potentially involving both Pol III-and Pol II-dependent transcription, although the contributions of the two polymerases require further validation.

In-depth studies in vertebrates have shown that *U6* promoter structure plays an important role in determining polymerase recruitment, with competition and dynamic equilibrium governing the recruitment of the two polymerases [43,44]. In the land plant *A. thaliana*, specific polymerase recruitment was found to be determined by the spacing between the USE element and TATA box in the promoter: a longer spacing recruits Pol II for U2–U5, whereas a shorter spacing recruits Pol III for U3 and U6 [67]. Nevertheless, the ability of *U6* promoters to drive *GUS* gene expression has also been demonstrated in several plants, suggesting that such dual transcriptional capability may be relatively widespread in plants [33,45]. Notably, *UpU6* promoters lacked USE elements yet supported both small RNA and protein-coding gene expression, suggesting that their transcriptional flexibility may be determined by unknown promoter architecture. Essentially, this dual small-RNA/mRNAmay arise from the structural similarity between type 3 Pol III promoters and Pol II promoters, as well as the shared regulatory factors between the Pol II and Pol III transcription systems [68,69]. Our results provided evidence suggesting that this dual transcriptional activity has already emerged in green algae lineage, rising the possibility that similar transcriptional properties are highly likely to be reproducible in other algae. Future studies will need to clarify the specific polymerases responsible for each type of transcription, as well as the potential mechanisms of selective recruitment.

Unexpectedly, the *UpU6* promoters were also demonstrated to be functional in driving *GUS* gene expression in *Agrobacterium*. To our knowledge, this represented the first report of *U6* promoter activity in a prokaryotic system. Previous studies have reported that some eukaryotic Pol II promoters are active in prokaryotes, which has been attributed to the significant similarity between their TATA box sequences and the bacterial -10 region (TATAAT) [70–72]. Given that eukaryotic transcription machinery evolved from Archaea and share a common ancestor with bacteria [73,74], this cross-system activity may reflect the structural conservation of primitive transcriptional mechanisms.

### 4.3 Editing efficiency and editing events

The gene editing results showed that *UpAPT* mutants were obtained in every experimental group, indicating that all three *UpU6* promoters were applicable to the vector-based approach and could effectively drive sgRNA transcription. The editing frequency calculated from the starting cell number ranged from 1.88 × 10^-6^ to 4.07 × 10^-6^, with no significant difference among the three *UpU6* promoters (Table 1). These values were comparable to those reported for simplex genome editing in *U. compressa* (7 × 10^-8^ to 3 × 10^-6^) [23], yet with reduced inter-group variation. On the other hand, the mutation-positive fraction among resistant thalli was 92.3% (n = 52), which was comparable to that reported for *U. compressa* (74.0%, n = 173) and could essentially meet the requirements for gene-editing experiments in *Ulva*. This suggested that the SISR process in *Ulva* facilitated chromatin opening, thereby promoting Cas9-mediated cleavage.

Notably, the three *UpU6* promoters exhibited distinct mutation spectra. Only substitutions and small indels were detected in the *UpU6-2* promoter group, whereas the *UpU6-1* and *UpU6-3* promoter groups displayed more diverse editing events, with large SVs also identified. To explain this difference, we proposed a hypothesis here, which was that it might be related to the variations in sgRNA expression levels driven by different promoters. Previous studies have shown that higher or more sustained sgRNA expression can elevate intracellular levels of active Cas9-sgRNA complexes, thereby increasing the frequency of target cleavage and opportunities for repeated cleavage [25,26]. Persistent or multiple DSB events can influence the extent of DNA end resection, which in turn affects the repair process and ultimate outcome of DSB repair [75,76]. To verify this hypothesis, future studies will quantify the abundance of sgRNAs and increase the sample size to meet the statistical requirements. Furthermore, it remains to be verified whether the effect of the promoter on the editing outcome is related to the specific target gene and sgRNA.

The results also showed that substitutions occurred at a high frequency of 45.5% (15/33) in CRISPR-Cas9-mediated gene editing in *U. prolifera*, which was consistent with previous reports in this species (16.7%) but markedly higher than those in other animals and plants, where small indels typically predominate [20,77]. This may reflect a distinct preference for DSB repair in *U. prolifera*. Additionally, a high proportion of SVs were detected, indicating that editing outcomes in *Ulva* were considerably complex. Further studies are needed to better understand the mechanisms underlying these outcomes.

### 4.4 Transposon insertion associated with gene editing

It should be noted that in this study, large fragment insertion events within the target gene were detected at a frequency of 18.2% (6/33) among the independent CRISPR-Cas9 edited mutants of *U. prolifera*. The length of the inserted sequence was 3933 bp, and the sequence was completely identical in multiple independent transformants. Additionally, one mutant was found where the 1150-bp insertion actually belonged to a part of the 3933-bp fragment. Sequence analysis revealed that the 3933-bp fragment had typical ∼111-bp TIRs at both ends, and an 8-bp TSD at the insertion site, suggesting that it belonged to the DNA transposon of the MULE superfamily [78]. Since the 3933-bp insertions were only detected within the *UpAPT* gene in experimental groups, and no corresponding mutation was detected in the negative control group, these findings suggested an association between MULE insertion and CRISPR-Cas9-mediated editing within the *UpAPT* gene. Studies have demonstrated that DNA damage-induced cellular stress can activate certain transposable elements, with transposition events preferentially occurring in chromatin-accessible regions [79,80]. In particular, the presence of TSD provided further support for the involvement of transposase. Therefore, we hypothesize that DSBs induced by CRISPR-Cas9 may increase chromatin accessibility at the target site, thereby creating a favorable integration environment for endogenous MULEs to transpose at the cleavage sites, although this proposed mechanism requires further validation.

Based on their transposition mechanisms and sequence structures, transposable elements are broadly divided into two major classes: retrotransposons and DNA transposons [81]. Transposable element sequences have been detected in the target genes of editing events in *Chlamydomonas* [82] and animals [83,84]. However, the insertion events reported in these reports all pertained to retrotransposons. To our knowledge, this study provided the first observation of insertions of endogenous DNA transposon at sites subjected to CRISPR-Cas9-mediated gene editing. Notably, no coexistence of transposition and conventional DSB repair outcomes was observed in the mutants, which raised the possibility of mutual exclusivity between these two processes, although further verification was required.

Transposons are ubiquitous components of eukaryotic genomes and serve as a major source of genomic variation and evolutionary innovation [85]. In *Ulva*, transposon sequences account for approximately 25.18% of the genome; however, their detailed composition remains to be fully characterized [86]. The MULE superfamily belongs to the DNA transposons and is distinguished by its exceptionally high mutagenic potential [87]. Members of this superfamily have been identified across fungi, plants, and animals [88]. Here, we identified a MULE-like transposable element family in *Ulva*. BLASTN analysis of the 3933-bp MULEsequence identified hundreds of hits in the *U. prolifera* genome, among which 10 non-redundant loci exhibited 99.85%–100% sequence identity across full-length alignments. A tBLASTn search using the transposase sequence within this MULE against the *U. compressa* genome also returned a significant hit. Given that MULE sequences are highly variable and may show pronounced interspecific divergence even within the same genus [89,90], their widespread distribution throughout *Ulva* cannot be excluded. Additionally, MULE-associated insertions were observed in up to 18.2% independent gene-editing events in this study, suggesting that this MULE family may retain a high transpositional activity in *Ulva*. These findings broadened our understanding of MULE distribution among green algae. The potential effects of MULEs on gene-editing efficiency and mutation spectrum in *Ulva*, as well as their influence on mutant stability, warrant further investigation.

### 4.5 Prospects for UpU6 promoter applications

This study revealed two key findings in *U. prolifera*: the dual small-RNA/mRNA transcriptional activity and cross-system functionality of the *UpU6* promoters, as well as the frequent occurrence of MULE insertions at CRISPR-Cas9 target sites. Together, these provided an important molecular basis for optimizing and developing gene editing tools in *Ulva*.

Three *UpU6* promoters have been confirmed as effective elements for gene editing, enriching the toolkit of type 3 Pol III promoters for future research on genome editing and RNAi in *U. prolifera*. Compared to existing RNP-based approaches [20–22], the vector-based CRISPR-Cas9 system established in this study would provide a more flexible platform than for development of multiplex genome editing in *U. prolifera*. Given their ability to support both small RNA transcription and protein-coding gene expression, a single *UpU6* promoter may potentially be employed to simultaneously express sgRNA and Cas9 when vector size reduction is needed to facilitate transformation, analogous to the validated application of the human H1 promoter [91]. Conversely, on the premise of having clarified the selective recruitment mechanism of the *UpU6* promoter, targeted mutations could be introduced into its upstream regulatory elements. This would engineer *UpU6* promoter variants with enhanced Pol II or Pol III activity and specificity for different transcriptional requirements [91,92].In recent years, new gene editing strategies such as transposase-assisted target site integration (TATSI) have been developed, which enable targeted knock-in of large DNA sequences carried by transposons. The insertion efficiency was significantly higher than that of the traditional HDR pathway [93]. In this study, frequent MULE insertions were identified at CRISPR-Cas9-targeted loci in *U. prolifera*. This observation raised the possibility that endogenous MULE elements may be explored as potential components of transposon-assisted genome engineering strategies, including large-fragment knock-in. Further analysis of the transposase activity and transposition mechanism of MULEs in *Ulva* is still needed to evaluate the feasibility of the above applications.

## Credit Author Statement

**Qingyun Hou:** Methodology, Validation, Investigation, Formal analysis, Visualization, Data curation; Writing -original draft. **Zhenghua Wang:** Resources, Methodology. **Jin Zhao:** Resources. **Peng Jiang:** Conceptualization, Supervision, Writing - review & editing, Project administration, Funding acquisition.

## Acknowledgements

This research was funded by the Southwest and Beibu Gulf Scientific Expedition Phase I Program (2026XNKK0600), Natural Science Foundation of Qingdao City (23-2-1-172-zyyd-jch), Shandong Provincial Natural Science Foundation (ZR2024MD028), and Key R&D Program of Shandong Province (2019GSF107012).

## Statement regarding informed consent, human/animal rights

No conflicts, informed consent, or human or animal rights are applicable to this study.

## Abbreviations

2-FA: 2-fluoroadenine
AtU6: *U6* snRNA gene of *Arabidopsis thaliana*
CrU6: *U6* snRNA gene of *Chlamydomonas reinhardtii*
DSB: double-strand break
G: guanine
indel: insertion-deletion mutation
ML: maximum likelihood
MUG: 4-methylumbelliferyl β-D-glucuronide
MULE: Mutator-like element
Pol II: RNA polymerase II
Pol III: RNA polymerase III
sgRNA: single-guide RNA
SISR: somatic *in situ* regeneration
SNP: single nucleotide polymorphism
snRNA: small nuclear RNA
SV: structural variation
T: thymine
TIR: terminal inverted repeat
TSD: target site duplication
TSS: transcription start site
UpAPT: *APT* gene of *Ulva prolifera*
UpU6: *U6* snRNA gene of *Ulva prolifera*
USE: upstream sequence element
VSE medium: Von Stosch’s Enriched medium
YSGT: the Yellow Sea green tide

## References

[1] R.L. Fletcher, The occurrence of “green tides”— a review, in: W. Schramm, P.H. Nienhuis (Eds.), Marine Benthic Vegetation, Springer Berlin Heidelberg, Berlin, Heidelberg, 1996: pp. 7–43, 10.1007/978-3-642-61398-2_2.

[2] Y. Zhan, Z. Qiu, Y. Wang, Y. Su, Y. Li, Y. Cui, S. Qu, P. Wang, X. Rong, Long-term spatiotemporal characteristics of *Ulva prolifera* green tide and effects of environmental drivers on its monitoring by satellites: A case study in the Yellow Sea, China, from 2008 to 2023, J. Mar. Sci. Eng. 12 (2024) 630, 10.3390/jmse12040630.

[3] Y. Xiao, J. Zhang, T. Cui, J. Gong, R. Liu, X. Chen, X. Liang, Remote sensing estimation of the biomass of floating *Ulva prolifera* and analysis of the main factors driving the interannual variability of the biomass in the Yellow Sea, Mar. Pollut. Bull. 140 (2019) 330–340, 10.1016/j.marpolbul.2019.01.037.

[4] Y. Zhang, P. He, H. Li, G. Li, J. Liu, F. Jiao, J. Zhang, Y. Huo, X. Shi, R. Su, N. Ye, D. Liu, R. Yu, Z. Wang, M. Zhou, N. Jiao, *Ulva prolifera* green-tide outbreaks and their environmental impact in the Yellow Sea, China, Natl. Sci. Rev. 6 (2019) 825–838, 10.1093/nsr/nwz026.

[5] M. Sun, Y. Li, Y. Ren, Y. Chen, Redefine sustainable fisheries targets under the impact of the Southern Yellow Sea green tide: mitigating the recurring surge in natural mortality, Front. Mar. Sci. 9 (2022) 813024, 10.3389/fmars.2022.813024.

[6] J. Zhao, P. Jiang, S. Qin, X. Liu, Z. Liu, H. Lin, F. Li, H. Chen, C. Wu, Genetic analyses of floating *Ulva prolifera* in the Yellow Sea suggest a unique ecotype, Estuar. Coast. Shelf Sci. 163 (2015) 96–102, 10.1016/j.ecss.2015.05.027.

[7] J. Cui, A.P. Monotilla, W. Zhu, Y. Takano, S. Shimada, K. Ichihara, T. Matsui, P. He, M. Hiraoka, Taxonomic reassessment of *Ulva prolifera* (Ulvophyceae, Chlorophyta) based on specimens from the type locality and Yellow Sea green tides, Phycologia 57 (2018) 692–704, 10.2216/17-139.1.

[8] Y. He, S. Shen, D. Yu, Y. Wang, J. Yin, Z. Wang, Y. Ye, The *Ulva prolifera* genome reveals the mechanism of green tides, J. Ocean. Limnol. 39 (2021) 1458–1470, 10.1007/s00343-020-0212-5.

[9] K. Tamura, H. Bono, Genome sequence of the edible green alga *Ulva prolifera*, originating from the Yoshinogawa River in Japan, Microbiol. Resour. Announc. 11 (2022), 10.1128/mra.00430-22.

[10] K. Jahan, M.S.A. Supty, J.S. Lee, K.H. Choi, Transcriptomic analysis provides new insights into the tolerance mechanisms of green macroalgae *Ulva prolifera* to high temperature and light stress, Biology 13 (2024) 725, 10.3390/biology13090725.

[11] H. Liu, T. Wu, J. Zhang, P. He, Exploring the over-wintering and over-summering mechanisms of *Ulva prolifera* from physiological and transcriptome perspectives and their impacts on green tides, Mar. Environ. Res. 210 (2025) 107310, 10.1016/j.marenvres.2025.107310.

[12] H. Gu, M. Li, J. Liu, H. He, X. Yang, S. Shen, Y. He, Regulation of *LOV1* in *Ulva prolifera* thermotolerance: A preliminary study, Plant Physiol. Biochem. 229 (2025) 110689, 10.1016/j.plaphy.2025.110689.

[13] M. Jinek, K. Chylinski, I. Fonfara, M. Hauer, J.A. Doudna, E. Charpentier, A programmable dual-RNA–guided DNA endonuclease in adaptive bacterial immunity, Science 337 (2012) 816–821, 10.1126/science.1225829.

[14] P.D. Hsu, E.S. Lander, F. Zhang, Development and applications of CRISPR-Cas9 for genome engineering, Cell 157 (2014) 1262– 1278, 10.1016/j.cell.2014.05.010.

[15] R. Suzuki, T. Yamazaki, A. Toyoda, S. Kawano, A transformation system using *rbcS* N-terminal region fused with GFP demonstrates pyrenoid targeting of the small subunit of RubisCO in *Ulva compressa*, Cytologia 79 (2014) 427–428, 10.1508/cytologia.79.427.

[16] W. Oertel, T. Wichard, A. Weissgerber, Transformation of *Ulva mutabilis* (Chlorophyta) by vector plasmids integrating into the genome, J. Phycol. 51 (2015) 963–979, 10.1111/jpy.12336.

[17] M. Kakinuma, M. Ikeda, D.A. Coury, H. Tominaga, I. Kobayashi, H. Amano, Isolation and characterization of the *rbcS* genes from a sterile mutant of *Ulva pertusa* (Ulvales, Chlorophyta) and transient gene expression using the *rbcS* gene promoter, Fish. Sci. 75 (2009) 1015–1028, 10.1007/s12562-009-0101-5.

[18] C. Wu, P. Jiang, Y. Guo, J. Liu, J. Zhao, H. Fu, Isolation and characterization of *Ulva prolifera actin1* gene and function verification of the 5′ flanking region as a strong promoter, Bioengineered 9 (2018) 124–133, 10.1080/21655979.2017.1325041.

[19] C. Wu, Establishment of stable expression system for exogenous gene in Ulva prolifera, Ph.D. thesis, Institute of Oceanology, Chinese Academy of Sciences, Qingdao, China, 2017.

[20] K. Ichihara, T. Yamazaki, S. Kawano, Genome editing using a DNA-free clustered regularly interspaced short palindromic repeats-Cas9 system in green seaweed *Ulva prolifera*, Phycol. Res. 70 (2022) 50–56, 10.1111/pre.12472.

[21] K. Ichihara, T. Yamazaki, S. Kawano, Cas9 ribonucleoproteins and single-strand oligodeoxynucleotides enable targeted insertional mutagenesis in *Ulva*, Algal Res. 81 (2024) 103598, 10.1016/j.algal.2024.103598.

[22] Z. Qin, W. Surnido, H. Mizuta, T. Uji, Stable transgene expression and CRISPR-mediated knock-in system of a bacteria-derived antibiotic selection gene in the green alga *Ulva prolifera*, BMC Plant Biol. 25 (2025) 1323, 10.1186/s12870-025-07411-y.

[23] J. Blomme, J. Arraiza Ribera, O. De Clerck, T.B. Jacobs, Consolidating *Ulva* functional genomics: gene editing and new selection systems, New Phytol. 246 (2025) 1710–1723, 10.1111/nph.70068.

[24] D. Kummari, S.R. Palakolanu, P.B.K. Kishor, P. Bhatnagar-Mathur, P. Singam, V. Vadez, K.K. Sharma, An update and perspectives on the use of promoters in plant genetic engineering, J. Biosci. 45 (2020) 119, 10.1007/s12038-020-00087-6.

[25] G. Yuen, F.J. Khan, S. Gao, J.M. Stommel, E. Batchelor, X. Wu, J. Luo, CRISPR/Cas9-mediated gene knockout is insensitive to target copy number but is dependent on guide RNA potency and Cas9/sgRNA threshold expression level, Nucleic Acids Res. 45 (2017) 12039–12053, 10.1093/nar/gkx843.

[26] N. Javaid, S. Choi, CRISPR/Cas system and factors affecting its precision and efficiency, Front. Cell Dev. Biol. 9 (2021) 761709, 10.3389/fcell.2021.761709.

[27] G.R. Kunkel, R.L. Maser, J.P. Calvet, T. Pederson, U6 small nuclear RNA is transcribed by RNA polymerase III, Proc. Natl. Acad. Sci. U.S.A. 83 (1986) 8575–8579, 10.1073/pnas.83.22.8575.

[28] F. Waibel, W. Filipowicz, U6 snRNA genes of *Arabidopsis* are transcribed by RNA polymerase Ⅲ but contain the same two upstream promoter elements as RNA polymerase Ⅱ-transcribed U-snRNA genes, Nucleic Acids Res. 18 (1990) 3451–3458, 10.1093/nar/18.12.3451

[29] S.D. Kor, N. Chowdhury, A.K. Keot, K. Yogendra, C. Chikkaputtaiah, P. Sudhakar Reddy, RNA Pol III promoters—key players in precisely targeted plant genome editing, Front. Genet. 13 (2023) 989199, 10.3389/fgene.2022.989199.

[30] F. Li, M. Xue, D. Guo, L. Zhu, Y. Li, L. Xie, A truncated endogenous U6 promoter enables high-efficiency CRISPR editing in flax (*Linum usitatissimum* L.), Plants 14 (2025) 1142, 10.3390/plants14071142.

[31] A. Bao, D.J. Burritt, H. Chen, X. Zhou, D. Cao, L.S.P. Tran, The CRISPR/Cas9 system and its applications in crop genome editing, Crit. Rev. Biotechnol. 39 (2019) 321–336, 10.1080/07388551.2018.1554621.

[32] X. Sun, Z. Hu, R. Chen, Q. Jiang, G. Song, H. Zhang, Y. Xi, Targeted mutagenesis in soybean using the CRISPR-Cas9 system, Sci. Rep. 5 (2015) 10342, 10.1038/srep10342.

[33] L. Long, D.D. Guo, W. Gao, W.W. Yang, L.P. Hou, X.N. Ma, Y.C. Miao, J.R. Botella, C.P. Song, Optimization of CRISPR/Cas9 genome editing in cotton by improved sgRNA expression, Plant Methods 14 (2018) 85, 10.1186/s13007-018-0353-0.

[34] A. Charrier, E. Vergne, N. Dousset, A. Richer, A. Petiteau, E. Chevreau, Efficient targeted mutagenesis in apple and first time edition of pear using the CRISPR-Cas9 system, Front. Plant Sci. 10 (2019) 40, 10.3389/fpls.2019.00040.

[35] Y.-S. Riu, G.H. Kim, K.W. Chung, S.G. Kong, Enhancement of the CRISPR/Cas9-based genome editing system in lettuce (*Lactuca sativa* L.) using the endogenous U6 promoter, Plants 12 (2023) 878, 10.3390/plants12040878.

[36] A.M. Domitrovich, G.R. Kunkel, Multiple, dispersed human U6 small nuclear RNA genes with varied transcriptional efficiencies, Nucleic Acids Res. 31 (2003) 2344–2352, 10.1093/nar/gkg331.

[37] A.L. Didychuk, S.E. Butcher, D.A. Brow, The life of U6 small nuclear RNA, from cradle to grave, RNA 24 (2018) 437–460, 10.1261/rna.065136.117.

[38] X. Qi, L. Dong, C. Liu, L. Mao, F. Liu, X. Zhang, B. Cheng, C. Xie, Systematic identification of endogenous RNA polymerase III promoters for efficient RNA guide-based genome editing technologies in maize, Crop J. 6 (2018) 314–320, 10.1016/j.cj.2018.02.005.

[39] Y.H. Di, X.J. Sun, Z. Hu, Q.Y. Jiang, G.H. Song, B. Zhang, S.S. Zhao, H. Zhang, Enhancing the CRISPR/Cas9 system based on multiple GmU6 promoters in soybean, Biochem. Biophys. Res. Commun. 519 (2019) 819–823, 10.1016/j.bbrc.2019.09.074.

[40] X. Xia, S. Li, N. Wang, P. Cheng, B. Zhu, P. Zhang, D. Yang, H. Lin, L. Niu, Convenient, high-efficiency multiplex genome editing in autotetraploid alfalfa using endogenous U6 promoters and visual reporters, aBIOTECH 6 (2025) 81–90, 10.1007/s42994-025-00200-z.

[41] X. Ma, Q. Zhu, Y. Chen, Y.G. Liu, CRISPR/Cas9 platforms for genome editing in plants: Developments and applications, Mol. Plant 9 (2016) 961–974, 10.1016/j.molp.2016.04.009.

[42] C. Ren, Y. Liu, Y. Guo, W. Duan, P. Fan, S. Li, Z. Liang, Optimizing the CRISPR/Cas9 system for genome editing in grape by using grape promoters, Hortic. Res. 8 (2021) 52, 10.1038/s41438-021-00489-z.

[43] J. Park, G.R. Kunkel, Both RNA polymerase III and RNA polymerase II accurately initiate transcription from a human U6 promoter in *vitro*, Biochem. Biophys. Res. Commun. 214 (1995) 934–940, 10.1006/bbrc.1995.2376.

[44] Z. Gao, E. Herrera-Carrillo, B. Berkhout, RNA polymerase II activity of type 3 Pol III promoters, Mol. Ther. Nucleic Acids 12 (2018) 135–145. 10.1016/j.omtn.2018.05.001.

[45] M. Kanai, K. Hikino, S. Mano, Cloning and functional verification of endogenous U6 promoters for the establishment of efficient CRISPR/Cas9-based genome editing in castor (*Ricinus communis*), Genes 14 (2023) 1327, 10.3390/genes14071327.

[46] S. Gao, M. Zhao, S. Sun, X. Fan, J. Yan, Y. Xin, Y. Zhan, F. Zeng, Development of an endogenous promoter-driven CRISPR/Cas9 system for genome editing in *Fraxinus mandshurica*, For. Res. 5 (2025) 0–0, 10.48130/forres-0025-0016.

[47] S. Riesenberg, P. Kanis, R. Karlic, T. Maricic, Robust prediction of synthetic gRNA activity and cryptic DNA repair by disentangling cellular CRISPR cleavage outcomes, Nat. Commun. 16 (2025) 4717, 10.1038/s41467-025-59947-0.

[48] P. Jiang, J. Wang, Y. Cui, Y. Li, H. Lin, S. Qin, Molecular phylogenetic analysis of attached Ulvaceae species and free-floating *Enteromorpha* from Qingdao coasts in 2007, Chin. J. Ocean. Limnol. 26 (2008) 276–279, 10.1007/s00343-008-0276-0.

[49] E. Leskinen, P. Pamilo, Evolution of the ITS sequences of ribosomal DNA in *Enteromorpha* (Chlorophyceae), Hereditas 126 (1997) 17–23, 10.1111/j.1601-5223.1997.00017.x.

[50] S. Shimada, N. Yokoyama, S. Arai, M. Hiraoka, Phylogeography of the genus *Ulva* (Ulvophyceae, Chlorophyta), with special reference to the Japanese freshwater and brackish taxa, J. Appl. Phycol. 20 (2008) 979–989, 10.1007/s10811-007-9296-y.

[51] Q. Liu, C. Wu, W. Xie, J. Zhao, P. Jiang, Validation of mating type-related markers in *Ulva prolifera* (Ulvophyceae, Chlorophyta) and their detection during various reproductive modes, Algal Res. 62 (2022) 102611, 10.1016/j.algal.2021.102611.

[52] M. Lescot, PlantCARE, a database of plant cis-acting regulatory elements and a portal to tools for in silico analysis of promoter sequences, Nucleic Acids Res. 30 (2002) 325–327, 10.1093/nar/30.1.325.

[53] R.A. Jefferson, S.M. Burgess, D. Hirsh, β-Glucuronidase from *Escherichia coli* as a gene-fusion marker, Proc. Natl. Acad. Sci. U.S.A. 83 (1986) 8447–8451, 10.1073/pnas.83.22.8447.

[54] W. Jiang, A.J. Brueggeman, K.M. Horken, T.M. Plucinak, D.P. Weeks, Successful transient expression of Cas9 and single guide RNA genes in *Chlamydomonas reinhardtii*, Eukaryot. Cell 13 (2014) 1465–1469, 10.1128/EC.00213-14.

[55] A.Q. Lin, Q.Y. Hou, J. Zhao, P. Jiang, Transcriptomic analysis of somatic in situ regeneration in green macroalga *Ulva prolifera* (Ulvophyceae, Chlorophyta), J. Mar. Sci. Eng. 14 (2026) 1705, 10.3390/jmse14181705.

[56] T. Martin, R. Wöhner, S. Hummel, L. Willmitzer, W. Frommer, The GUS gene reporter system as a tool to study plant gene expression, in: S. Gallagher (Ed.), GUS Protocols: Using the GUS Gene as a Reporter of Gene Expression, Academic Press, San Diego, 1992: pp. 23–43.

[57] H. Liu, Y. Ding, Y. Zhou, W. Jin, K. Xie, L.L. Chen, CRISPR-P 2.0: An improved CRISPR-Cas9 tool for genome editing in plants, Mol. Plant 10 (2017) 530–532, 10.1016/j.molp.2017.01.003.

[58] S. Connelly, Small nuclear RNA genes transcribed by either RNA polymerase II or RNA polymerase III in monocot plants share three promoter elements and use a strategy to regulate gene expression different from that used by their dicot plant counterparts, Mol. Cell. Biol. 14 (1994) 5910–5919.

[59] D.A. Brow, C. Guthrie, Spliceosomal RNA U6 is remarkably conserved from yeast to mammals, Nature 334 (1988) 213–218, 10.1038/334213a0.

[60] R. Reddy, D. Henning, G. Das, M. Harless, D. Wright, The capped U6 small nuclear RNA is transcribed by RNA polymerase III, J. Biol. Chem. 262 (1987) 75–81, 10.1016/S0021-9258(19)75890-6.

[61] G. Das, D. Henning, D. Wright, R. Reddy, Upstream regulatory elements are necessary and sufficient for transcription of a U6 RNA gene by RNA polymerase III, EMBO J. 7 (1988) 503–512, 10.1002/j.1460-2075.1988.tb02838.x.

[62] F. Solymosy, T. Pollák, Uridylate-rich small nuclear RNAs (UsnRNAs), their genes and pseudogenes, and UsnRNPs in plants: Structure and function. A comparative approach, Crit. Rev. Plant Sci. 12 (1993) 275–369, 10.1080/07352689309701904.

[63] F. Guerineau, R. Waugh, The U6 small nuclear RNA gene family of potato, Plant Mol. Biol. 22 (1993) 807–818, 10.1007/BF00027367.

[64] B.B. Wang, V. Brendel, The ASRG database: Identification and survey of *Arabidopsis thaliana* genes involved in pre-mRNA splicing, Genome Biol. 5 (2004) R102, 10.1186/gb-2004-5-12-r102.

[65] K.E. Giles, M. Caputi, K.L. Beemon, Packaging and reverse transcription of snRNAs by retroviruses may generate pseudogenes, RNA 10 (2004) 299–307, 10.1261/rna.2150604.

[66] L. Schramm, N. Hernandez, Recruitment of RNA polymerase III to its target promoters, Genes Dev. 16 (2002) 2593–2620, 10.1101/gad.1018902.

[67] F. Waibel, W. Filipowicz, RNA-polymerase specificity of transcription of *Arabidopsis* U snRNA genes determined by promoter element spacing, Nature 346 (1990) 199–202, 10.1038/346199a0.

[68] D. Raha, Z. Wang, Z. Moqtaderi, L. Wu, G. Zhong, M. Gerstein, K. Struhl, M. Snyder, Close association of RNA polymerase II and many transcription factors with Pol III genes, Proc. Natl. Acad. Sci. U.S.A. 107 (2010) 3639–3644, 10.1073/pnas.0911315106.

[69] S.Z. Shah, T.N. Perry, A. Graziadei, V. Cecatiello, T. Kaliyappan, A.D. Misiaszek, C.W. Müller, E.P. Ramsay, A. Vannini, Structural insights into distinct mechanisms of RNA polymerase II and III recruitment to snRNA promoters, Nat. Commun. 16 (2025) 141, 10.1038/s41467-024-55553-8.

[70] T.K. Antonucci, P. Wen, W.J. Rutter, Eukaryotic promoters drive gene expression in *Escherichia coli*, J. Biol. Chem. 264 (1989) 17656–17659, 10.1016/S0021-9258(19)84621-5.

[71] D. Jacob, A. Lewin, B. Meister, B. Appel, Plant-specific promoter sequences carry elements that are recognised by the eubacterial transcription machinery, Transgenic Res. 11 (2002) 291–303, 10.1023/A:1015620016472

[72] S. Goussard, C. Grillot-Courvalin, P. Courvalin, Eukaryotic promoters can direct protein synthesis in Gram-negative bacteria, Microb. Physiol. 6 (2003) 211–218, 10.1159/000077252.

[73] L. Eme, A. Spang, J. Lombard, C.W. Stairs, T.J.G. Ettema, Archaea and the origin of eukaryotes, Nat. Rev. Microbiol. 15 (2017) 711–723, 10.1038/nrmicro.2017.133.

[74] O.V. Bylino, A.N. Ibragimov, Y.V. Shidlovskii, Evolution of regulated transcription, Cells 9 (2020) 1675, 10.3390/cells9071675.

[75] C. Xue, E.C. Greene, DNA repair pathway choices in CRISPR-Cas9-mediated genome editing, Trends Genet. 37 (2021) 639–656, 10.1016/j.tig.2021.02.008.

[76] M. Kosicki, F. Allen, F. Steward, K. Tomberg, Y. Pan, A. Bradley, Cas9-induced large deletions and small indels are controlled in a convergent fashion, Nat. Commun. 13 (2022) 3422, 10.1038/s41467-022-30480-8.

[77] J. De Saeger, E. Coulembier Vandelannoote, H. Lee, J. Park, J. Blomme, Genome editing in macroalgae: Advances and challenges, Front. Genome Ed. 6 (2024), 10.3389/fgeed.2024.1380682.

[78] A.A. Ferguson, N. Jiang, Mutator-Like elements with multiple long terminal inverted repeats in plants, Comp. Funct. Genomics 2012 (2012) 1–14, 10.1155/2012/695827.

[79] E.A. Farkash, E.T.L. Prak, DNA damage and L1 retrotransposition, BioMed Res. Int. 2006 (2006) 037285, 10.1155/JBB/2006/37285.

[80] Z. Wang, R. Schwacke, R. Kunze, DNA damage-induced transcription of transposable elements and long non-coding RNAs in *Arabidopsis* is rare and ATM-dependent, Mol. Plant 9 (2016) 1142–1155, 10.1016/j.molp.2016.04.015.

[81] D.J. Finnegan, Eukaryotic transposable elements and genome evolution, Trends Genet. 5 (1989) 103–107, 10.1016/0168-9525(89)90039-5.

[82] D. Guzmán-Zapata, J.M. Sandoval-Vargas, K.S. Macedo-Osorio, E. Salgado-Manjarrez, J.L. Castrejón-Flores, M.D.C. Oliver-Salvador, N.V. Durán-Figueroa, F. Nogué, J.A. Badillo-Corona, Efficient editing of the nuclear *APT* reporter gene in *Chlamydomonas reinhardtii* via expression of a CRISPR-Cas9 module, Int. J. Mol. Sci. 20 (2019) 1247, 10.3390/ijms20051247.

[83] R. Ono, M. Ishii, Y. Fujihara, M. Kitazawa, T. Usami, T. Kaneko-Ishino, J. Kanno, M. Ikawa, F. Ishino, Double strand break repair by capture of retrotransposon sequences and reverse-transcribed spliced mRNA sequences in mouse zygotes, Sci. Rep. 5 (2015) 12281, 10.1038/srep12281.

[84] J. Tao, Q. Wang, C. Mendez-Dorantes, K.H. Burns, R. Chiarle, Frequency and mechanisms of LINE-1 retrotransposon insertions at CRISPR/Cas9 sites, Nat. Commun. 13 (2022) 3685, 10.1038/s41467-022-31322-3.

[85] D. Lisch, How important are transposons for plant evolution, Nat. Rev. Genet. 14 (2013) 49–61. 10.1038/nrg3374.

[86] P. Garcia-Jimenez, R.R. Robaina, Exploring transposons in macroalgae: LTR elements and neighboring genes in red seaweeds, Front. Mar. Sci. 12 (2025) 1592442, 10.3389/fmars.2025.1592442.

[87] D. Lisch, Mutator transposons, Trends Plant Sci. 7 (2002) 498–504, 10.1016/S1360-1385(02)02347-6.

[88] M. Dupeyron, K.S. Singh, C. Bass, A. Hayward, Evolution of *Mutator* transposable elements across eukaryotic diversity, Mobile DNA 10 (2019) 12, 10.1186/s13100-019-0153-8.

[89] Z. Yu, S.I. Wright, T.E. Bureau, Mutator-like elements in *Arabidopsis thaliana*: Structure, diversity and evolution, Genetics 156 (2000) 2019–2031, 10.1093/genetics/156.4.2019.

[90] M. Nuruzzaman, T. Kanno, R. Amada, Y. Habu, I. Kasajima, T. Ishikawa, M. Kawai-Yamada, H. Uchimiya, Does the upstream region possessing MULE-like sequence in rice upregulate *PsbS1* gene expression? PLoS ONE 9 (2014) e102742, 10.1371/journal.pone.0102742.

[91] Z. Gao, E. Herrera-Carrillo, B. Berkhout, A single H1 promoter can drive both guide RNA and endonuclease expression in the CRISPR-Cas9 system, Mol. Ther. Nucleic Acids 14 (2019) 32–40, 10.1016/j.omtn.2018.10.016.

[92] Z. Gao, Y.U. Van Der Velden, M. Fan, C.A. Van Der Linden, M. Vink, E. Herrera-Carrillo, B. Berkhout, Engineered miniature H1 promoters with dedicated RNA polymerase II or III activity, J. Biol. Chem. 296 (2021) 100026, 10.1074/jbc.RA120.015386.

[93] P. Liu, K. Panda, S.A. Edwards, R. Swanson, H. Yi, P. Pandesha, Y.H. Hung, G. Klaas, X. Ye, M.V. Collins, K.N. Renken, L.A. Gilbertson, V. Veena, C.N. Hancock, R.K. Slotkin, Transposase-assisted target-site integration for efficient plant genome engineering, Nature 631 (2024) 593–600, 10.1038/s41586-024-07613-8.

